# Robustness of mitochondrial biogenesis and respiration explain aerobic glycolysis

**DOI:** 10.1101/2024.07.04.601975

**Authors:** Easun Arunachalam, Felix C. Keber, Richard C. Law, Chirag K. Kumar, Yihui Shen, Junyoung O. Park, Martin Wühr, Daniel J. Needleman

## Abstract

A long-standing observation is that in fast-growing cells, respiration rate declines with increasing growth rate and is compensated by an increase in fermentation, despite respiration being more efficient than fermentation. This apparent preference for fermentation even in the presence of oxygen is known as aerobic glycolysis, and occurs in bacteria, yeast, and cancer cells. Considerable work has focused on understanding the potential benefits that might justify this seemingly wasteful metabolic strategy, but its mechanistic basis remains unclear. Here we show that aerobic glycolysis results from the saturation of mitochondrial respiration and the decoupling of mitochondrial biogenesis from the production of other cellular components. Respiration rate is insensitive to acute perturbations of cellular energetic demands or nutrient supplies, and is explained simply by the amount of mitochondria per cell. Mitochondria accumulate at a nearly constant rate across different growth conditions, resulting in mitochondrial amount being largely determined by cell division time. In contrast, glucose uptake rate is not saturated, and is accurately predicted by the abundances and affinities of glucose transporters. Combining these models of glucose uptake and respiration provides a quantitative, mechanistic explanation for aerobic glycolysis. The robustness of specific respiration rate and mitochondrial biogenesis, paired with the flexibility of other bioenergetic and biosynthetic fluxes, may play a broad role in shaping eukaryotic cell metabolism.

## Introduction

In eukaryotes, the cellular energy currency adenosine triphosphate (ATP) is primarily generated by mitochondrial respiration^1^. In this process, the electron transport chain (ETC) uses electrons derived from the oxidation of carbon sources to reduce oxygen. The biochemical basis of respiratory flux control has been intensely studied^2–7^, but despite many important insights, key questions regarding the control of oxygen consumption under physiological conditions are still unanswered^8–11^. Notably, the mechanism that underlies the variation in respiration rate with growth rate is unclear^9,12–16^. In both prokaryotes and eukaryotes, it has been observed that as growth rates increase, the rate of respiration decreases, and cells begin to ferment even in the presence of oxygen – a phenomenon known variously as aerobic glycolysis, overflow metabolism, the Crabtree effect, and the Warburg effect^17–23^. This behavior is counterintuitive because fermentation is approximately an order of magnitude less ATP-efficient than respiration^1^.

In this work, we investigated respiratory flux control in the budding yeast *Saccharomyces cerevisiae*. We found that acute perturbations of ATP-consuming processes and acute alteration of nutrient supply did not affect respiration rate. However, extended cultivation in different carbon sources led to differences in respiration rate which could be explained by differences in mitochondrial content. We show that both the observed homeostasis of respiration rate given a fixed amount of mitochondria and the scaling of respiration with mitochondrial volume are due to the saturation of the electron transport chain.

To understand what underpins differences in mitochondrial content, and hence differences in respiration rate, we used live-cell imaging to measure the rate of mitochondrial biogenesis. We found that the rate at which mitochondrial mass accumulates remains similar across different growth conditions, even as cell division times vary considerably. When cell division times are longer, there is more time for mitochondria to accumulate, and thus the average amount of mitochondria per cell increases. Our findings lead to a saturation-accumulation-division (SAD) model of respiratory flux control: the ETC is saturated, mitochondria accumulate at a similar rate under different growth conditions, and mean mitochondrial amount is thus determined largely by division time. Combining the SAD model with a model of glucose uptake, based on the kinetics of glucose transporters and the external glucose concentration, quantitatively predicts the increase in fermentation and the decrease in respiration with increasing external glucose levels. The SAD model thus explains how the saturation of mitochondrial respiration and the robustness of mitochondrial biogenesis together give rise to aerobic glycolysis.

## Results

### Respiration rate remains constant regardless of changing ATP consumption or nutrient availability

Given that the oxidation of carbon sources and the consumption of ATP are coupled to oxygen consumption, we sought to understand the extent to which respiration rate is set by ATP demand or nutrient supply. To test the extent to which ATP demand controls oxygen consumption rate (OCR), we acutely perturbed the rates of ATP-consuming processes in ethanol-grown cells, which rely exclusively on respiration to produce ATP (Fig. 1A). We performed these experiments within minutes to characterize the *in situ* biochemical properties of the mitochondrial machinery already present, rather than changes in respiration rate that might result from adaptation of the proteome. We inhibited processes which previous work suggests are significant ATP consumers^24–27^: we decreased the rate of translation (using anisomycin), inhibited microtubule assembly (using nocodazole) and actin polymerization (using Latrunculin A), and altered ion pumping (using high salt). These perturbations of ATP demand had the expected phenotypic effects (Fig. S1A-H) and significantly impacted growth rate and cellular ATP levels (Fig. 1B-C), but did not significantly affect cellular oxygen consumption rate (Fig. 1D). Thus, while perturbing key ATP-consuming processes affects overall cellular metabolic state, it does not significantly affect respiration rate.

**Figure 1:**
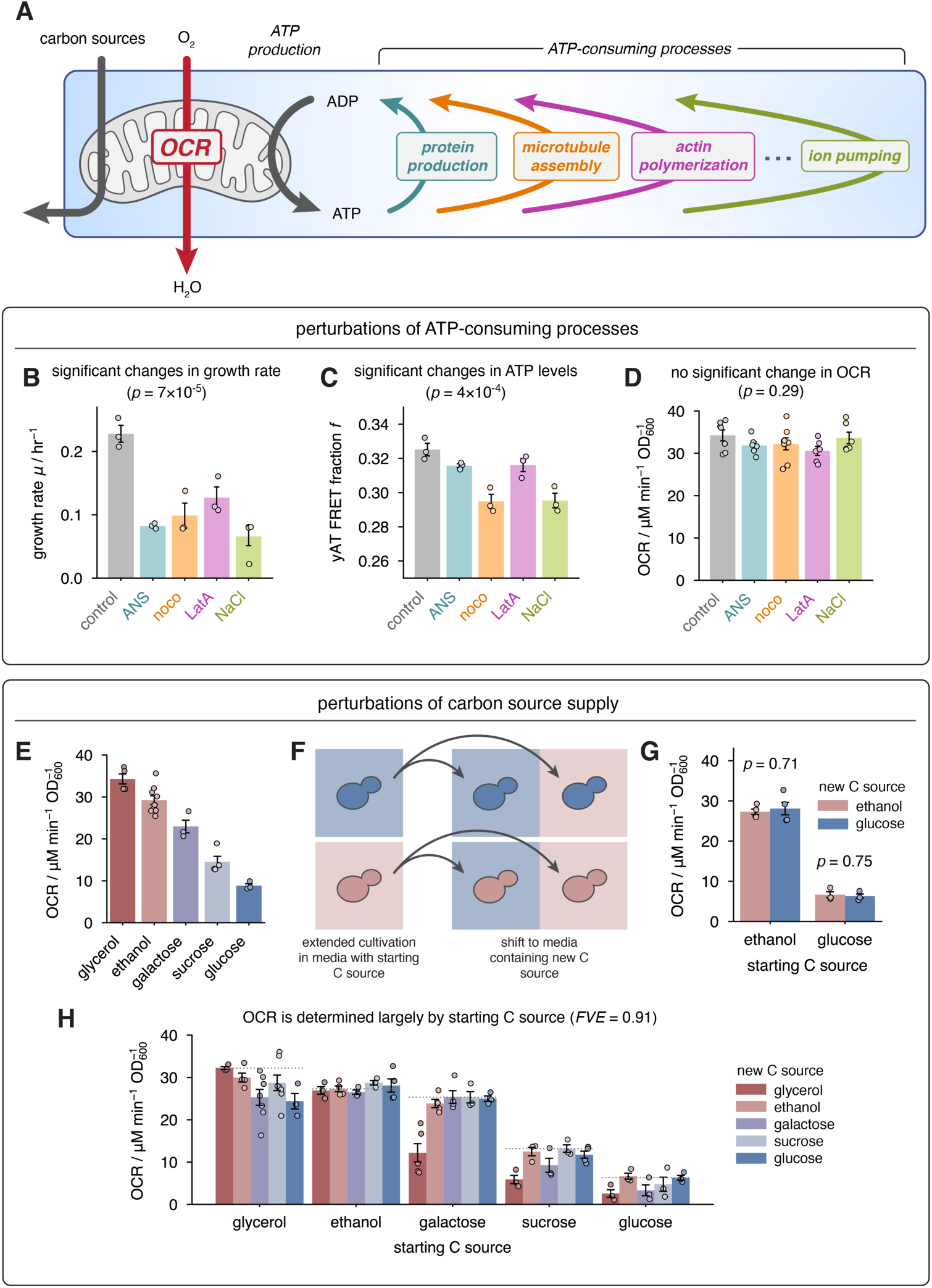
Respiration rate is insensitive to physiological perturbations of ATP demand and nutrient supply. **(A)** Mitochondria couple carbon source catabolism, oxygen consumption, and ATP production; many different cellular processes consume ATP. Perturbing the rate of different ATP consuming processes **(B)** causes significant changes to growth rate (*n* = 3 biological replicates), and **(C)** causes significant changes to ATP concentrations, as assayed by the FRET sensor yAT1.03 (*n* = 3 biological replicates), but **(D)** does not significantly affect OCR (*n* ≥ 6 biological replicates). In **(B)**-**(D)** all data are shown as mean ± s.e.m., and *p*-values are obtained by a one-way ANOVA. **(E)** Extended growth in different carbon sources results in different OCR (*n* ≥ 3 biological replicates). **(F)** Cells grown for an extended period in one carbon source were shifted to another carbon source before metabolic measurements. **(G)** OCR remains unchanged following shifts between glucose and ethanol (*n* ≥ 3 biological replicates). *p*-values are obtained by Welch’s two-tailed *t*-test. **(H)** OCR following pairwise shifts between five different carbon sources is largely explained by the preshift carbon source (*n* ≥ 3 biological replicates). Dotted reference lines indicate OCR for shifts where the starting and ending media are the same (no change in carbon source). *FVE* = fraction of variance in mean OCR of different conditions that is explained by initial carbon source. In **(E)**, **(G)**, and **(H)** all data are shown as mean ± s.e.m.

We next asked whether the external carbon supply determines the rate of respiration. It is well-known that extended growth in different carbon sources results in different rates of oxygen consumption (Fig. 1E and^6,28,29^). However, these differences are accompanied by changes in growth rate, mode of metabolism, carbohydrate and lipid composition, and proteome composition^30–32^. Hence, measurements of cells well-adapted to growth on different carbon sources cannot establish whether the external carbon supply at the moment of the measurement determines respiration rate. To address this, we cultivated cells in media containing either ethanol or glucose, then shifted them to media containing the other carbon source immediately before measuring respiration rate (Fig. 1F). Because the media switch and measurement occurred within minutes, factors including the proteome (i.e. metabolic enzyme content), macromolecular stores, and mitochondrial content remained approximately constant. To determine if energy metabolism was impacted, we used fluorescence lifetime imaging microscopy to measure intracellular levels of the key redox coenzymes NAD(P)H, and found they change upon carbon source shifts (Fig. S1I-J). However, respiration rate remains unchanged: cells grown on ethanol retain the same high OCR when transferred to glucose-containing media as when transferred to ethanol-containing media, and glucose-grown cells have a low OCR when transferred to either glucose- or ethanol-containing media (Fig. 1G). We measured OCR following nutrient shifts and found that it adapted over several hours, on the timescale of growth and division (Fig. S2A-B).

To evaluate the generality of this phenomenon, we grew cells in media containing one of five different carbon sources (glycerol, ethanol, galactose, sucrose, or glucose), rapidly shifted them to new media containing each of the carbon sources in turn, and measured their OCR. We observed the same general trend: the carbon source in which cells were grown for an extended period before the shift explained the vast majority of variation (91%) in post-shift respiration rates (Fig. 1H). Overall, these data (Fig. 1) demonstrate that acute perturbations of either energetic demand or carbon supply only minimally affect OCR, suggesting that respiration is saturated.

### Respiration scales with mitochondrial content

We next sought to determine why cells grown in different carbon sources exhibit different respiration rates. We hypothesized that differences in mitochondrial content might contribute to this variation. Using confocal microscopy, we imaged mitochondrial networks in individual cells to reconstruct their 3-dimensional structure^33^ and calculate their volume (Fig. 2A). We measured the mean mitochondrial volume per cell and respiration rate under 17 conditions (the five carbon sources studied above as well as six glucose-limited cultures with and without amino acids). We observed a strong linear relationship between the rate of oxygen consumption per cell and the mean mitochondrial volume per cell (*R*^2^ = 0.89; Fig. 2B). Each 1 µm^3^-increase in mitochondrial volume resulted in the same ~15 µM min^−1^ OD^−1^change in OCR, consistent with the hypothesis that each additional unit of mitochondrial volume contains the same metabolic enzymes, which operate at the same rate. To test the relationship between mitochondrial volume and metabolic enzyme content, we performed multiplexed proteomics of cells grown under each of the 17 conditions. We found that the total abundance of mitochondrial proteins was proportional to mean single-cell mitochondrial network volume (*R*^2^ = 0.71 and *p* = 0.52, two-tailed *t*-test of the null hypothesis that the *y*-intercept is zero; Fig. 2C). Mitochondrial volume was highly correlated with the abundance of different groups of respiration-related enzymes (median *R*^2^ = 0.71; Fig. S3), indicating a constant addition of these enzymes per unit increase in mitochondrial volume.

**Figure 2:**
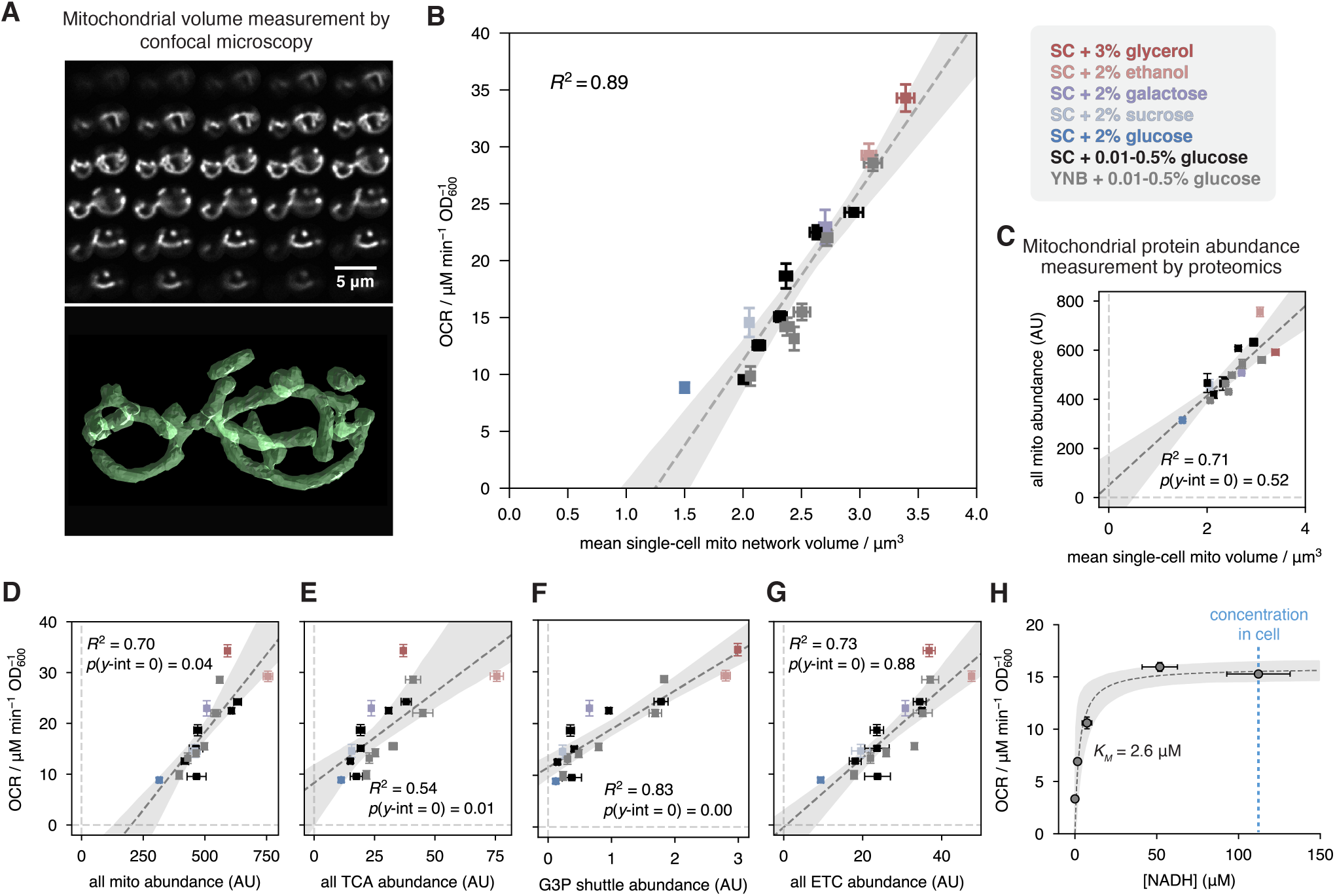
Mitochondrial content explains OCR differences because of electron transport chain saturation by NADH. **(A)** Confocal micrographs of mitochondria-targeted mNeonGreen (0.2 µm-spaced *z*-slices) were used to reconstruct the 3-dimensional structure of mitochondrial networks in single cells. **(B)** Mitochondrial network volume is linearly related to OCR across a variety of carbon sources. Points are shown as mean ± s.e.m. of *n* ≥ 208 cells per condition across *n* ≥ 3 biological replicates. SC = synthetic complete media, YNB = yeast nitrogen base media. **(C)** Mean single-cell mitochondrial volume and total abundance of mitochondrial proteins (as measured by proteomics, *n* = 3 biological replicates) are proportional. Dashed line represents linear regression of protein abundance against mitochondrial volume. **(D)** Linear regression of OCR against abundances of different functional classes of respiration-related proteins reveals that, while OCR is correlated with the abundance of various protein groups, it is strictly proportional only to electron transport chain (ETC) content. In **(C)** and **(D)** all data are shown as mean ± s.e.m., and *p*-values, estimated by bootstrapping, indicate the probability of observing the associated *y*-intercept, or one more extreme, given the null hypothesis that the *y*-intercept is zero. **(E)** OCR measurements and NADH concentration measurements for cells treated with different concentrations of IAA are consistent with saturation of the ETC by NADH. Data are shown as mean ± s.e.m. (*n* ≥ 3 biological replicates). In **(B)**-**(E)** dashed lines indicate best fit from regressions, and shaded regions indicate the 95% confidence interval from bootstrapping.

### Protein and metabolite levels are consistent with electron transport chain saturation by NADH

To determine which mitochondrial enzymes are saturated and thus control respiration rate, we investigated how the abundance of different functional groups of enzymes scaled with respiration rate. We reasoned that if OCR is controlled by a given functional group, then the amount of that group should not just be linearly related to OCR, but strictly proportional to OCR (such that OCR is zero in the absence of that group). We therefore performed linear regressions of OCR against the abundance of candidate groups, and identified those groups that had a nearly proportional relationship (i.e. a small absolute value of the regression *y*-intercept) and significant explanatory power (high *R*^2^) (Fig. S4A-B). For each regression, we tested the null hypothesis that the *y*-intercept was zero (i.e. not proportional) by bootstrap sampling.

We examined the trend of respiration with respect to total abundance of mitochondrial proteins (Fig. 2D), and found a strong linear relationship, consistent with previous work^9^, but with a large negative *y*-intercept (*p* = 0.04), indicating that the mitochondrial proteome as a whole is not proportional to OCR. We next considered the enzymes of the tricarboxylic acid (TCA) cycle, which generate reducing equivalents that are used for respiration; OCR is not proportional to these enzymes either (*p* = 0.01, Fig. 2E). Recent work has suggested that in cancer cells, the malate-aspartate shuttle and the glycerol-3-phosphate shuttle set the flux of cytosolic reducing equivalents into mitochondria^16^. However, the fit of OCR against the abundance of shuttles in our data suggests that they do not control OCR in yeast (*p* < 0.01, Fig. 2F; and S4C). We further tested this hypothesis by knocking out the matrix-facing NADH dehydrogenase *NDI1*, required for oxidation of NADH transported into mitochondria by shuttles, and *GUT2*, a component of the glycerol-3-phosphate shuttle. Consistent with previous work^5,34^, we found that OCR did not change significantly (Fig. S5A). This result suggests that shuttles are not the bottleneck for respiration, though it does not rule out the possibility of compensatory re-wiring of redox metabolism in response to these knockouts. Finally, we considered the total abundance of electron transport chain (ETC) complexes, which we found to be proportional to OCR (*p* = 0.88; Fig. 2G). Many individual components of the ETC are similarly proportional to OCR (Fig. S4C).

If ETC abundance controls respiration rate, we would expect primary electron acceptors to be saturated by the species which donate electrons, which are primarily reduced nicotinamide adenine nucleotides (NADH). To test whether NADH is saturating, we investigated how OCR is impacted by alteration of NADH levels in live cells. We manipulated NADH levels in glucose-grown cells, which generate NADH via glyceraldehyde-3-phosphate (GAPDH), by titrating a GAPDH inhibitor (IAA). We compared OCR with NADH levels which we measured by mass spectrometry and by fluorescence lifetime imaging (Fig. 2E and Fig. S5B-E). We observed a relationship consistent with Michaelis-Menten kinetics, with a half-maximal rate obtained at *K*_M_ = 2.6 ± 1.4 µM. This is far lower than the physiological NADH concentration of 112.2 ± 4.8 µM, consistent with electron transport chain saturation by NADH. We find that respiration rate is similarly insensitive to perturbations of mitochondrial membrane potential: decreasing membrane potential with a protonophore did not increase respiration rate (Fig. S5F-H).

Taken together, our data indicate that the ETC is saturated by NADH, leading to an insensitivity to perturbations of ATP demand and nutrient supply, and that ETC content is linearly related to mitochondrial volume. Hence, differences in respiration rate across different growth conditions are largely due to differences in mitochondrial volume (and thus ETC content).

### Mitochondrial content is largely controlled by division time

We next asked how mitochondrial volume was controlled across the different carbon sources studied here. It has previously been proposed that nutrient supply-specific mitochondrial biogenesis controls the amount of mitochondria present under different growth conditions^35–38^.

To test whether this was the case, we directly measured mitochondrial biogenesis rates in individual cells under different conditions using time-lapse confocal imaging (Fig. 3A). The increase in mitochondrial network volume was approximately linear in time during both the G1 phase of the cell cycle and over the course of budding, albeit with different slopes (Fig. 3B and Fig. S6A-C). We measured the rate of mitochondrial biogenesis during G1 and budding in the five carbon sources studied earlier, and calculated cell-cycle-averaged rates 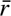 from estimates of G1 vs. budding fraction (Fig. 3C and Fig. S6D). Surprisingly, mitochondrial biogenesis rates did not vary substantially across different conditions (coefficient of variation CV ≈ 0.2), while the variation in cell cycle times was considerably greater (CV ≈ 0.5). Furthermore, mitochondrial biogenesis rates were only weakly correlated with mean mitochondrial volume (Pearson’s *r* = −0.40), while cell cycle times were strongly correlated (Pearson’s *r* = −0.80). We sought to understand the regulation of mitochondrial volume using a simple mathematical model (Supplementary Note 1). In this model of single-cell mitochondrial volume dynamics, mitochondrial volume accumulates continuously over the duration of the cell cycle. Thus, the average mitochondrial volume per cell is approximately proportional to both the average mitochondrial biogenesis rate 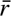 and the cell cycle time *T*. To test this model, we used it to predict the average mitochondrial volume per cell using parameters estimated from single-cell microscopy of mitochondrial networks and bulk doubling time measurements. The predicted mitochondrial volumes were in good agreement with direct measurements of mean volumes (Fig. 3E). These results support an “accumulation-division” model of mitochondrial volume control, in which mitochondria continually accumulate over the course of the cell cycle such that longer times between successive divisions provide more time for mitochondria to accumulate, and hence greater average mitochondrial volumes per cell (Fig. 3F).

**Figure 3:**
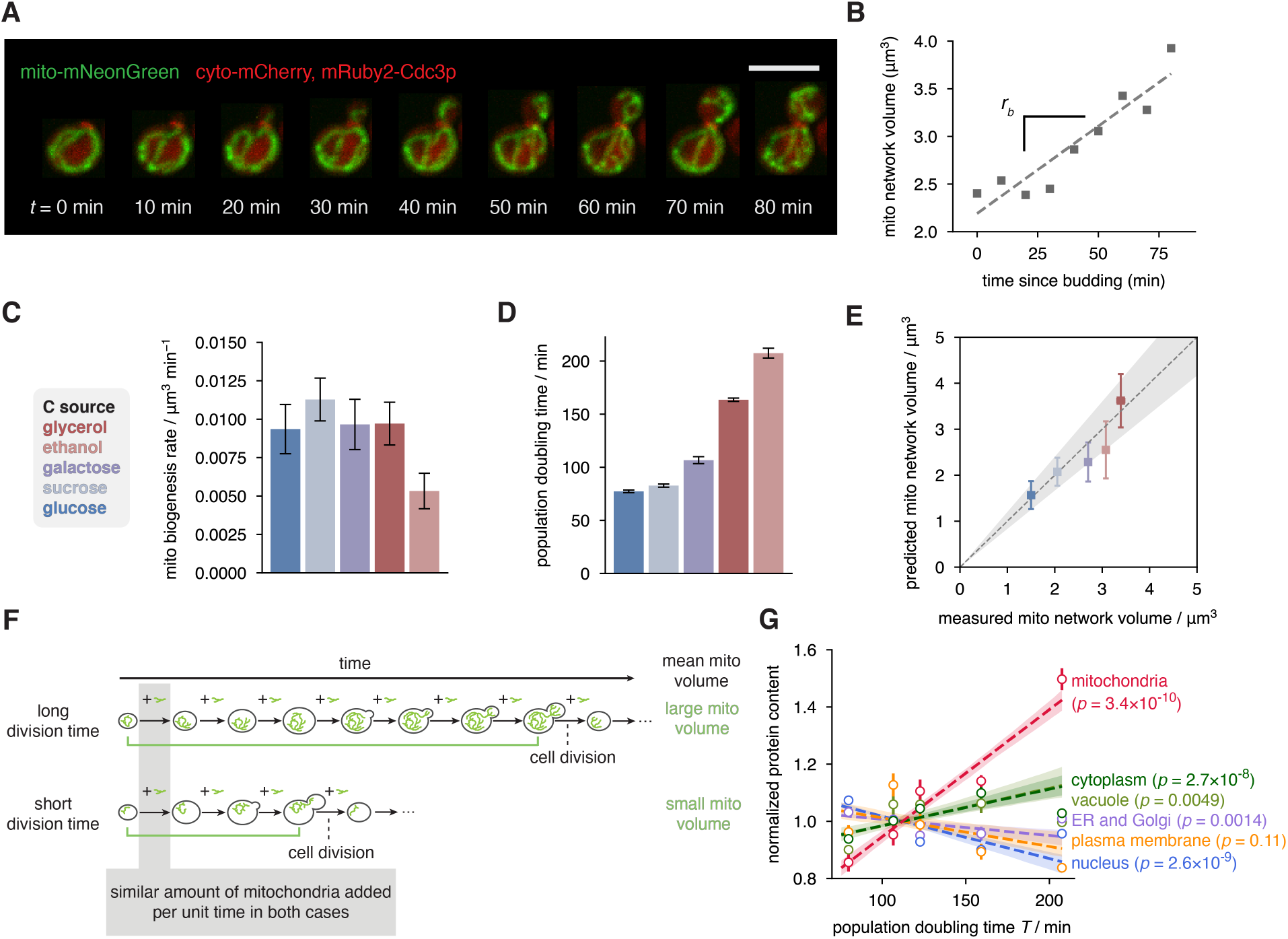
Variation in cell division time is the primary determinant of differences in mitochondrial content across various carbon sources. **(A)** Time-lapse confocal imaging of the mitochondrial network in a single cell. The max-*z* projections of 3d images are shown; scale bar is 5 µm. **(B)** Extraction of single-cell mitochondrial biogenesis rate from linear regression of mitochondrial network volume over time. **(C)** Cell-cycle-averaged mitochondrial biogenesis rate for cells cultivated in SC medium with different carbon sources. Data are shown as mean ± s.e.m. calculated by bootstrapping of *n* ≥ 151 mitochondrial growth trajectories across G1 and budding from *n* = 3 biological replicates. **(D)** Population doubling times in different carbon sources. Data are shown as mean ± s.e.m., *n* ≥ 4 biological replicates. **(E)** Predicted average mitochondrial volume per cell, calculated from measured mitochondrial biogenesis rates and cell cycle times (Supplementary Note 1) agree with mean measured mitochondrial network volumes. Points represent mean ± s.e.m. **(F)** Schematic of single-cell mitochondrial volume over the course of a cell cycle in conditions with long and short division times. **(G)** Total abundance of mitochondrial, ER and Golgi, cytoplasmic, vacuolar, membrane-associated, and nuclear proteins as a function of population doubling time. The 17 different growth conditions were grouped into five bins according to doubling time for clarity; the mean doubling time and organelle protein content for each of these bins is shown. Error bars indicate 68% confidence intervals from bootstrapping. *p*-values indicate the probability that each regression slope would be observed given the null hypothesis that organelle abundance is independent of doubling time.

Mitochondria are unique among organelles in that they maintain their own genome and gene expression machinery^39^; hence, we hypothesized that the accumulation-division model might apply specifically to them and not other organelles. To test this we investigated the extent to which the abundance of different organelles could be explained by differences in cell cycle times as predicted by the accumulation-division model. We quantified the total amount of protein in different organelles, including mitochondria, the nucleus, the endoplasmic reticulum and Golgi apparatus, as well as proteins in the cytoplasm and cell membrane, across different growth conditions. The normalized abundance of proteins in each of these non-mitochondrial locations was a weak function of division time (relative changes all < 20%, Fig. 3G). In contrast, the amount of mitochondrial proteins varies drastically (relative change = 59%): under the conditions studied here, the slowest-growing cells possess nearly twice as much mitochondrial protein as fast-growing cells.

Taken together, our results are consistent with a saturation-accumulation-division (SAD) model that explains trends in respiration rate in fast-growing cells: (i) in each cell, respiration rate is set by the amount of mitochondria because respiration-associated machinery is saturated; (ii) because mitochondria accumulate over the course of the cell cycle, mitochondrial amount per cell is largely determined by the time between successive divisions.

### The source of reducing equivalents for respiration is similar under fermenting and non-fermenting conditions

The SAD model provides a simple explanation of respiratory flux control in which oxygen consumption rate varies smoothly with cell division time. However, prior work has shown that above a critical growth rate, glucose-grown yeast undergo a transition in energy metabolism and switch from non-fermentative to fermentative growth (i.e. begin to perform aerobic glycolysis)^14,19^. To understand how these observations can be reconciled with one another, we sought to test if mitochondria themselves undergo a metabolic transition as growth rate changes.

We investigated the impact of growth rate on fluxes in central carbon metabolism using a series of glucose-limited cultures. The growth rates of these cultures increased monotonically with glucose concentration (Fig. 4A). We measured glucose consumption, ethanol production, and oxygen consumption rates, and, as expected, we observed ethanol production beyond the threshold glucose concentration of ~0.6 mM (Fig. 4B). To determine the contribution of different intramitochondrial fluxes to the observed respiration rate, we conducted parallel labeling experiments with 1,2-^13^C_2_-glucose or U-^13^C-glucose. We constructed a minimal model of glycolysis and the TCA cycle and performed ^13^C-metabolic flux analysis (MFA)^40^, which we constrained using metabolite labeling patterns and absolute extracellular fluxes (Fig. 4C). This analysis revealed two trends: firstly, the relative contribution of reducing equivalents produced by the TCA cycle is qualitatively similar across these conditions (Fig. 4D); secondly, full TCA cycle turning occurs under most of the glucose concentrations studied, and ceases only at the highest glucose concentrations (Fig. S7A-E). While ethanol production begins at a low glucose concentration (above ~0.6 mM), qualitative changes in TCA metabolism emerge only at a 10-fold higher glucose concentration (~5.6 mM). Thus, these tracing experiments show no evidence of a major transition in mitochondrial TCA cycle metabolism accompanying the onset of fermentation.

**Figure 4:**
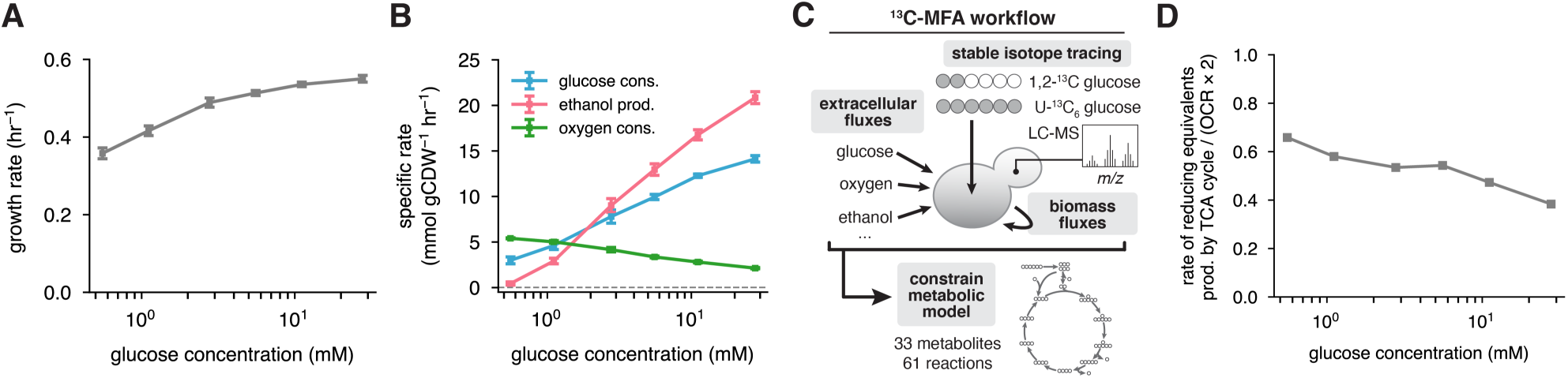
The transition to aerobic glycolysis is not driven by a transition in TCA cycle fluxes. **(A)**-**(B)** Growth rate, glucose consumption, ethanol production, and oxygen consumption rates in dilute glucose-limited batch cultures, as a function of glucose concentration in the media. Mean ± s.e.m. (*n* ≥ 6 biological replicates). **(C)** Extracellular flux measurements and stable isotope tracing were integrated into ^13^C -MFA, which enabled inference of fluxes through central carbon metabolism in each glucose-limited culture. **(D)** Contribution of TCA cycle-derived reducing equivalents to measured oxygen consumption rate.

### Glycolysis and fermentation are not saturated

To determine how yeast switch from non-fermentative to fermentative growth in the absence of a dramatic transition in mitochondrial metabolism, we next investigated the control of glycolysis and fermentation. Glycolysis produces ATP that supplies various energy-consuming cellular processes, and it produces the carbon and reducing equivalents consumed by fermentation and respiration. We inhibited microtubule polymerization (an energy-consuming process) using nocodazole or partially inhibited glycolysis using iodoacetic acid, and measured extracellular fluxes and growth rate. While OCR remained unchanged (Fig. 5A), growth rate, glucose consumption, and ethanol production all decreased dramatically (Fig. 5B-D). These acute perturbations indicate that unlike respiration, glycolysis and fermentation are not saturated: they can change in response to alterations of coupled fluxes^41^.

**Figure 5:**
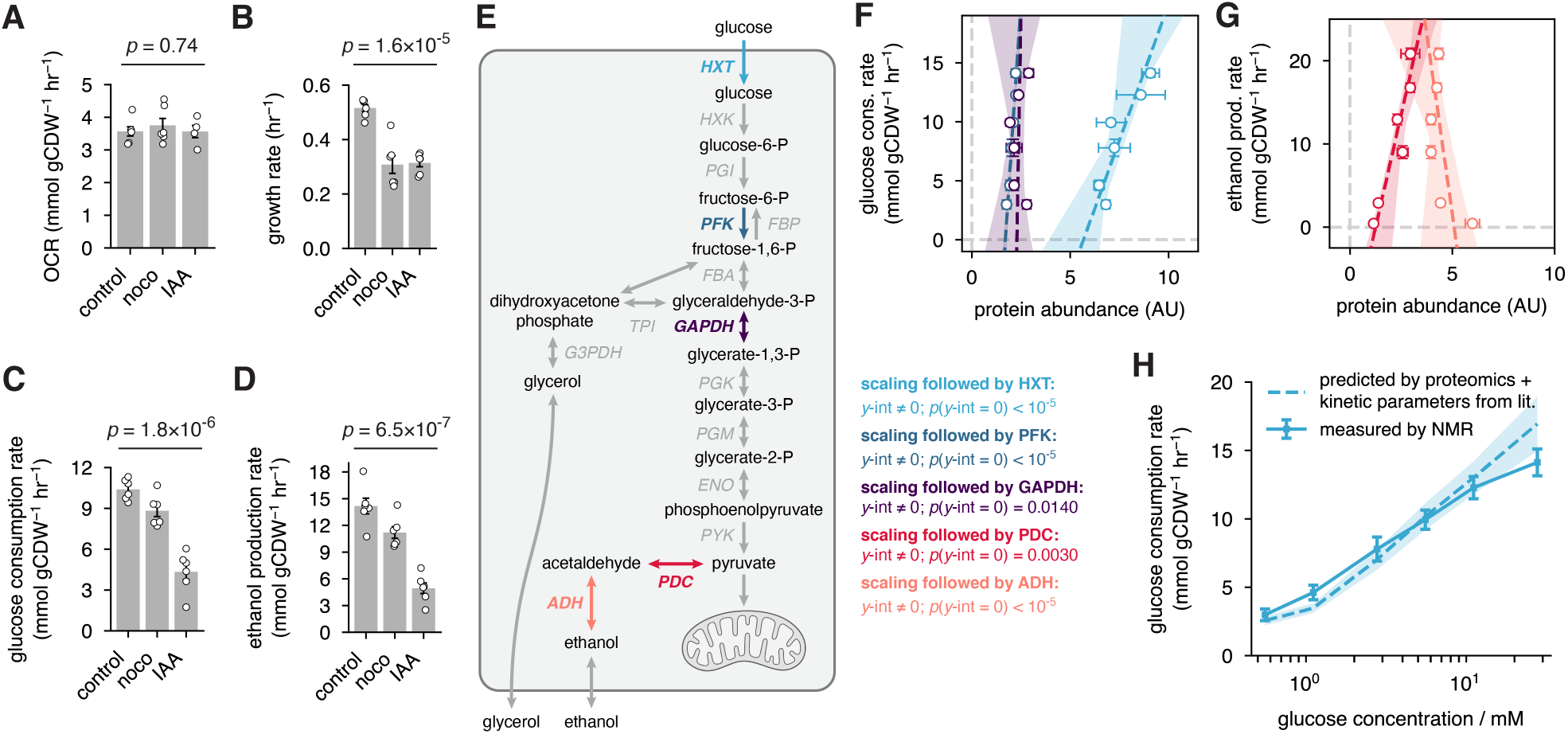
Glycolysis and fermentation are not saturated. **(A)-(D)** Acute inhibition of microtubule polymerization with nocodazole (noco), and partial inhibition of glycolysis with IAA, each **(A)** do not significantly perturb oxygen consumption (*n* ≥ 4 biological replicates), but **(B)-(D)** decrease growth, glucose consumption, and ethanol production (*n* = 6 biological replicates). Data are shown as mean ± s.e.m. **(E)** Key reactions in glycolysis and fermentation. The abundances of enzymes facilitating rate-determining reactions in **(F)** glycolysis and **(G)** fermentation are not proportional to the flux through those pathways (colored lines). Data are shown as mean ± s.e.m. (*n* = 3 biological replicates for proteomics, and *n* ≥ 6 for fluxes). Dashed lines indicate linear regressions of enzyme abundance against flux, and shaded regions indicate 95% CI for regressions. **(H)** Measured glucose consumption rate (solid line) is well-predicted by Michaelis-Menten kinetics.

To further investigate the control of these pathways, we examined the correlation between glucose consumption and ethanol production rates and the abundance of key enzymes (Fig. 5E), as measured by proteomics. Glucose consumption rate is not proportional (*y*-int ≠ 0) to the abundance of glycolytic enzymes, including hexose transporters (HXT), phosphofructokinase (PFK), or glyceraldehyde-3-phosphate dehydrogenases (GAPDH) (Fig. 5F). In the case of fermentation, neither pyruvate decarboxylases (PDC) nor alcohol dehydrogenases (ADH) are proportional to ethanol production rate (Fig. 5G). These results again indicate that, unlike respiration, fermentation is not saturated. This is consistent with previous reports that many metabolic reactions in yeast are primarily controlled by changes in metabolite levels^42^, implying that the enzymes which carry out those reactions are not saturated.

It has previously been argued that glucose uptake rate in yeast can be explained by external glucose concentration and the kinetics of glucose transporters^43^. We sought to test if this held true in our glucose-limited cultures as well. We used the abundance of the different hexose transporters (measured via proteomics) and the reported values of their Michaelis-Menten kinetic parameters (measured *in vitro*^44^) to predict glucose uptake rates in each glucose-limited culture. Our results were in good agreement with measured values (Fig. 5H and Supplementary Note 2). Thus, unlike respiration, in which the ETC is saturated by NADH, glycolysis is not saturated. Instead, the rate of glucose uptake is determined by external glucose levels, and the rate of fermentation is flexible.

### An integrated model of redox balance explains aerobic glycolysis

Glycolysis, respiration, and fermentation are linked through redox balance: glycolysis and the TCA cycle generate reducing equivalents, and these are consumed by respiration and fermentation (Fig. 6A). We sought to determine if our models of glucose uptake rate and respiration rate could thus predict the rate of fermentation.

**Figure 6:**
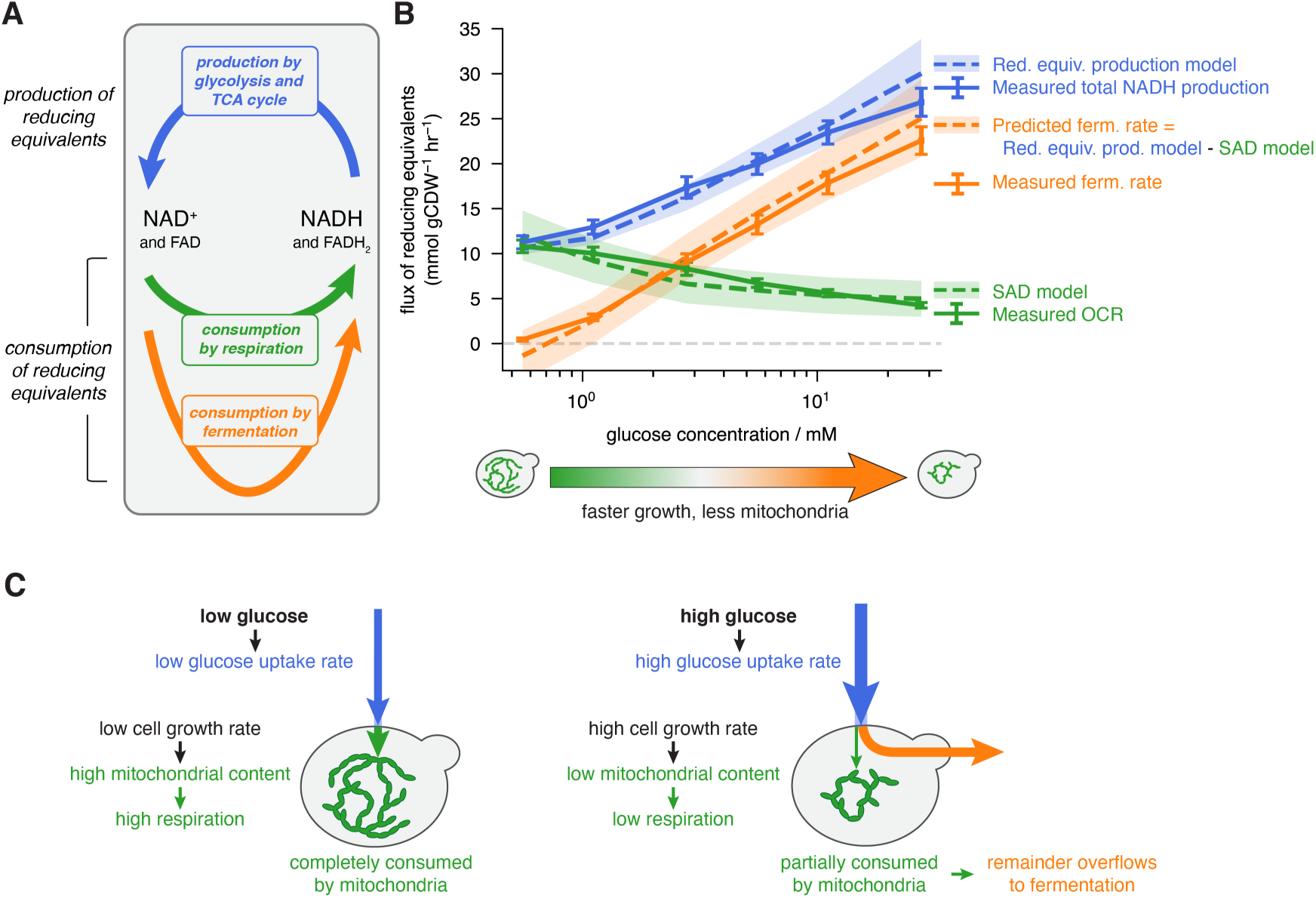
Aerobic glycolysis is explained by the SAD model and the kinetics of glucose transporters. **(A)** At steady state, production and consumption of reducing equivalents are balanced. **(B)** Flux of reducing equivalent production, and consumption by respiration and fermentation, as a function of glucose concentration. Ethanol production begins when reducing equivalent production exceeds consumption by respiration. Solid lines and error bars indicate mean ± s.e.m. for measurements; dashed lines and shaded regions indicate mean ± s.e.m. for model predictions. **(C)** Summary of physiological differences under low- and high-glucose conditions which underlie aerobic glycolysis.

We first calculated the total rate at which reducing equivalents were produced using a simple empirical model relating glucose uptake rate, glycolytic flux, and TCA cycle flux (Supplementary Note 3 and Fig. S8). The predictions of this model (Fig. 6B, dashed blue line) compared favorably with rates of reducing equivalent production determined directly from extracellular flux measurements (Fig. 6B, solid blue line). We next used the SAD model to calculate the rate at which respiration consumed reducing equivalents (Fig. 6B, dashed green line) which are in agreement with rates determined from direct OCR measurements (Fig. 6B, solid green line). At steady state, the total rate of production and consumption of reducing equivalents must be balanced. Thus, we calculated the fermentation rate by taking the difference between the total rate of reducing equivalent production (blue) and the rate of consumption by respiration (green). The predicted fermentation rate (Fig. 6B, dashed orange line) agrees with the measured fermentation rate (Fig. 6B, solid orange line).

At the lowest glucose concentration, there is no fermentation. As glucose concentration increases, glucose uptake rate increases, as does growth rate. The increase in growth rate leads to a decrease in mitochondrial content, and thus respiratory oxidative capacity. Increasing glycolytic flux and decreasing respiratory flux leads to a continual increase in fermentation with glucose concentration. Therefore, the combination of the glucose uptake rate model and the SAD model provide a mechanistic explanation for aerobic glycolysis (Fig. 6C).

## Discussion

Here we have combined respirometry, quantitative imaging, proteomics, and stable isotope tracing to identify trends in mitochondrial growth and metabolism across a range of conditions in budding yeast. These analyses have led to the saturation-accumulation-dilution (SAD) model of respiratory flux control: respiration operates at a constant rate per unit mitochondria, and hence the amount of mitochondria per cell determines cellular respiration rate; mitochondrial biogenesis occurs at a nearly constant rate across growth conditions, and hence the amount of mitochondria per cell is set largely by cell division time. The SAD model provides a quantitative, mechanistic explanation for respiratory flux control and aerobic glycolysis in budding yeast.

Previous work has focused on identifying potential fitness benefits associated with fermentation under aerobic conditions^21–23^. In contrast, the present study provides a mechanistic explanation of aerobic glycolysis: that this phenomenon arises naturally from the combination of the SAD model of respiration and the kinetics of glucose transporters. This finding reinforces the idea that the onset of fermentation is driven by the production of reducing equivalents exceeding respiratory capacity^12,45^.

Our work highlights three avenues for future work on mitochondrial metabolism and biogenesis. First, because respiration drives ATP production, it seems counterintuitive that inhibiting processes that consume ATP can leave respiration unchanged. However, it is conceivable that cells possess ATP flux-buffering mechanisms that maintain total ATP consumption, or alternatively, that mitochondria may decouple respiration from ATP production by modulating proton leak^10^. Second, the mechanism underlying the similar rates of mitochondrial biogenesis across different conditions is not yet clear^9^. Numerous processes, including lipid synthesis, membrane assembly, and synthesis and import of proteins must all be coordinated to enable mitochondrial biogenesis^37,46–49^, but which of these is rate-determining is not understood. Furthermore, the mechanism limiting mitochondrial biogenesis may be different in different growth regimes, as suggested by the scaling of mitochondrial content with growth rate when cells divide very slowly^6,9,50^. Even in these alternative growth regimes, differential regulation of mitochondrial biogenesis rates and cell growth rates may play an important role in determining mitochondrial abundance, and hence regulating metabolic fluxes. Third, it is not yet clear to what extent the SAD model describes respiratory flux control and fermentation in other systems. Recent work has provided evidence that part of this model – that mitochondria are saturated by NADH – may be true in mammalian cells: respiration in mouse oocytes is similarly insensitive to perturbations of ATP demand and nutrient supply^8^, and it has been argued that mitochondria are saturated in cancer cells^16^. However, the role of membrane potential in controlling respiration rate may be more complex in these cells. It is also currently unknown if in other systems variation in mitochondrial amount is the primary driver of variation in respiration rate, and whether cell division time underlies the differences in mitochondrial amount.

Though the molecular players involved in central carbon metabolism are well-known and have been thoroughly characterized, it has remained unclear what sets fluxes through these pathways. Here, we have developed a phenomenological model of respiratory control and aerobic glycolysis. The quantitative, coarse-grained approach we have employed may guide future efforts to develop a systems-level understanding of other aspects of metabolism and growth.

## Methods

### Strains and culture conditions

All strains used in this study were prototrophic W303 derivatives (see Table S1 for a detailed list of strains and their genotypes.) Yeast were cultivated at 30°C with agitation in synthetic complete (SC) or yeast nitrogen base (YNB) media. The carbon sources used were 2% w/v glucose, 2% w/v sucrose, 2% w/v galactose, 2% v/v ethanol, and 3% v/v glycerol (for different carbon source experiments), and 0.5%, 0.2%, 0.1%, 0.05%, 0.02%, and 0.01% w/v glucose (for glucose limitation experiments). Cells were grown for at least 15 doubling periods and maintained in exponential phase for at least two doubling periods prior to all measurements. Growth rates were determined by measurements of optical density at 600 nm (OD_600_) using a Genesys 30 Visible spectrophotometer (Thermo Scientific). The correspondence between optical density and cell dry weight was determined by filtering exponential-phase cultures, drying them at 65°C for 24 hours, then measuring the mass and subtracting that of the dried filter.

### ATP demand, nutrient supply, and membrane potential perturbations

For ATP demand perturbations, imaging, growth rate, and respirometry measurements were performed on cells cultivated in SC medium with 2% v/v ethanol, which produce ATP exclusively by respiration.

For protein synthesis inhibition experiments, cells were treated with 300 µM anisomycin (ANS) for 30 min before measurements. Action was verified by measuring nascent protein synthesis using the Click-iT kit (Click Chemistry Tools) as per manufacturer recommendations. In brief, cells were treated with or without ANS, then transferred to methionine-free SC medium supplemented with 500 µM L-homopropargylglycine (HPG), with or without ANS, for 40 minutes. Following HPG incorporation, cells were fixed with 4% paraformaldehyde in PBS, permeabilized with 0.5% Triton X-100 in PBS, then stained with Alexa Fluor 555 azide. Nascent protein was visualized by fluorescence microscopy.

For microtubule polymerization inhibition experiments, cells were treated with 66 µM nocodazole (noco) for 30 min before measurements. A strain with labeled tubulin (Venus-Tub1p), in which microtubule bundles are normally visible, was used to verify that nocodazole had dissolved bundles.

For actin polymerization inhibition experiments, cells were treated with 200 µM Latrunculin A (LatA) for 5 min, and then diluted into a larger volume for respirometry or imaging. We verified by microscopy that actin polymerization remained inhibited following dilution. A strain with a labeled actin-binding protein (Abp140p-mNeonGreen), in which the branched actin network is normally visible, was used to verify the dissolution of the network.

For high-salt perturbation experiments, cells were transferred to SC medium containing 2% v/v ethanol and 200 mM NaCl before measurements, which is known to increase ion pumping activity^1^. Because acute exposure to high sodium is known to decrease cytosolic pH^2^, we verified the physiological effect of NaCl treatment by measuring pH using the genetically encoded sensor pHluorin2^3,4^ using a fluorescence lifetime readout^5^.

For growth rate measurements, culture density was measured for three points following each inhibition, except in the case of LatA treatment, for which two points were used. ATP concentrations were measured using a genetically encoded Förster resonant energy transfer (FRET) biosensor, yAT1.03^6^. Changes in bound state were measured using fluorescence lifetime imaging, as described below. For nutrient shift experiments, exponentially growing cells were harvested by centrifugation, washed once in the new medium, then resuspended in the new medium. Measurements were typically completed within 30 minutes of the shift. For membrane potential perturbations, cells were treated with 200 nM carbonyl cyanide-*p*-trifluoromethoxyphenylhydrazone (FCCP) for 30 min before imaging or respirometry.

### Microscopy

#### Sample preparation

Glass-bottomed dishes were coated with a solution of 4 mg/mL concanavalin A type IV (Sigma-Aldrich) for 5 min, rinsed with culture media, filled with cell suspension for 5 min, rinsed using fresh media to remove unbound cells, and finally filled with 1 mL of media for imaging. Cells were maintained at 30°C using a stage-top heater box (Ibidi) and an objective heater (Bioptechs).

#### Fluorescence lifetime imaging

Fluorescence lifetime imaging microscopy (FLIM) of yAT1.03, NAD(P)H, and pHLuorin2 was performed with a two-photon laser scanning microscope controlled by LabVIEW (National Instruments). Excitation was provided by an Insight X3 tunable pulsed laser operating at 80 MHz (Spectra-Physics) and emission was detected using HPM-100-40 photomultiplier tubes and SPC-150 time-resolved single photon counting cards (Becker & Hickl). A 40x 1.2 NA water immersion objective (Nikon) was used for all measurements.

yAT1.03 imaging was performed with 865 nm excitation and a 482/35 emission filter; NAD(P)H was imaged using 750 nm excitation and a 460/50 emission filter; and pHluorin2 was imaged with 927 nm excitation and a 525/50 emission filter (all filters were purchased from Semrock). For yAT1.03 and NAD(P)H imaging, we fit normalized fluorescence lifetime decays *F* (*t*) to a two-exponential model (with signal amplitude *A*, long lifetime *τ*_*l*_, short lifetime *τ*_*s*_, and short lifetime fraction *f*), convolved with the instrument response function (IRF):

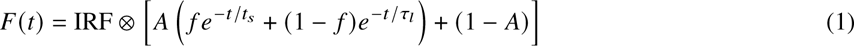

The instrument response function was measured using second harmonic generation from a urea crystal. Heatmaps of mean pHluorin2 fluorescence lifetimes were obtained by computing the mean arrival time for each cell pixel, averaged over neighboring cell pixels, weighted by a Gaussian kernel with a standard deviation of 30 pixels. Arrival times were corrected by subtracting the peak arrival time of the instrument response function.

#### Confocal imaging

Confocal imaging of mitochondrial networks was performed with a Nikon Eclipse Ti microscope equipped with a CSU-X1 spinning disk unit (Yokogawa), an ORCA Flash CMOS camera (Hamamatsu), 488 and 560 nm laser lines (Spectral Applied Physics), and a 60x 1.2 NA water immersion objective (Nikon).

Mitochondrial network structure was visualized using an mNeonGreen fluorescent protein targeted to the mitochondrial matrix using a pre-Su9 sequence (mito-mNeonGreen^7^). Mito-mNeonGreen was imaged using 488 nm excitation and a 525/50 nm emission filter. Cytoplasmic mCherry was used as a fiducial marker for segmentation of individual cells. A labeled septin ring component, mRuby2-Cdc3p, which is only visible when cells are not in G1, was used as a cell cycle marker^8,9^. Cytoplasmic mCherry and mRuby2-Cdc3p were simultaneously imaged using 560 nm excitation and a 594/30 nm emission filter. *z*-stacks were acquired with a spacing of 0.2 µm for mNeonGreen and 1 µm for mCherry/mRuby2 using MicroManager 1.4 controlled by a custom Beanshell script.

#### Measurement of mitochondrial volume and biogenesis rate

Automatic instance segmentation of cells was performed with CellPose^10^. Cells which could not be unambiguously segmented due to growth out of the field of view or overlapping in *z* were excluded from the segmentation. Time-lapse tracking of cell masks was performed with btrack^11^. Masks were manually corrected where necessary. Cell cycle trajectories were manually annotated on the basis of cell morphology and signal from mRuby2-Cdc3p.

Mitochondrial networks were segmented by Mitograph^12^ which produces a skeleton and surface mesh for each network. Segmented tubules were stretched in *z* relative to *xy* due to anisotropy in the point spread function, so the geometry of individual tubules was corrected by traversing the network skeleton and shrinking the distance to the nearest mesh point to ensure that the average profile was cylindrical. The volume enclosed by the corrected surface mesh was used for further analysis.

#### Membrane potential measurements

Cells were grown in SC+0.1% glucose to mid-exponential phase, then stained for 30 minutes with 100 nM tetramethyl-rhodamine (TMRM; Sigma-Aldrich). Cells were harvested by centrifugation, resuspended in TMRM-free media, and imaged by confocal microscopy. Mitochondria were segmented using Otsu thresholding.

### Oxygen consumption rate measurements

An oxygen-sensitive electrode (OX-50, Unisense A/S) was calibrated using air-saturated media and media sparged with nitrogen gas as endpoints. A chamber with the electrode was filled with cell suspension and then sealed. Oxygen depletion was monitored for 10-20 minutes, and the initial portion of the oxygen concentration trace, during which equilibration takes place, was discarded. A linear regression was performed to calculate the slope (i.e. the oxygen consumption rate), which was then normalized by cell density.

### Uptake and secretion rate measurements

Glucose consumption and ethanol, glycerol, and pyruvate production rates were determined using serial measurements of media composition in batch cultures, similar to previously described procedures^13^. Overnight cultures were back-diluted to OD 0.02-0.1 and grown for at least two doublings before beginning measurements to ensure cells were in the exponential growth phase. Cell density was measured and media aliquots were collected for at least three points during the early portion of the growth curve, when cultures were dilute and growing exponentially, to ensure that concentrations remained similar over the course of the experiment. For experiments involving acute perturbations, cells were treated with either 200 µM iodoacetic acid or 66 µM nocodazole for 20 minutes before beginning sampling.

A 3-(trimethylsilyl)-1-propanesulfonic acid (DSS-d_6_; 50 mM in D_2_O) internal standard was diluted 1:10 in spent medium, which was then analyzed by ^1^H NMR (400 MHz, Bruker). Spectra were collected using the *zgesgp* pulse sequence, and analyzed with MestReNova software. The following chemical shifts were used for quantitation: 0 ppm (s, 9H) for DSS-d_6_, 3.46 ppm (m) for glucose, and 1.17 ppm (t, 3H) for ethanol. Calibration curves based on standards of known glucose and ethanol content were used to calculate concentrations of these species in media samples. Glycerol and pyruvate content were quantified by LC-MS.

The ratio of extracellular flux to growth rate was determined by a linear fit of glucose, ethanol, glycerol, and pyruvate concentration against culture density over time. The growth rate under each condition was determined by a linear fit of the logarithm of cell density over time. The product of these two values yielded the absolute glucose, ethanol, glycerol, and pyruvate fluxes.

### Isotope tracing and LC-MS analysis

Cells were grown for at least 24 hours in SC medium with the appropriate concentration of glucose, then harvested by centrifugation, washed once in fresh medium containing the same concentration of ^13^C glucose, resuspended in fresh SC + ^13^C glucose at OD_600_ 0.05-0.1, and grown for 4-6 hours. Parallel tracing experiments were performed for each glucose concentration: one with 100% 1,2-^13^C glucose, and the other with 100% U-^13^C glucose. Exponentially growing cells were harvested by vacuum filtration on nylon membrane filters (0.45 µm) resting on a fritted glass support. Filters were quickly rinsed with 1 mL of yeast nitrogen base (YNB) media without glucose, and immediately flipped cell-side-down into 400 µL of -20°C 40:40:20 high-performance liquid chromatography (HPLC) grade acetonitrile:methanol:water in a six-well plate to rapidly quench metabolism. Extraction was continued for 20 min at -20°C, following which filters were flipped cell-side-up and thoroughly washed with the extraction solvent in the well. Metabolite extracts were collected in an Eppendorf tube, centrifuged at 4°C for 10 min, re-extracted with 100 µL of fresh solvent, and centrifuged once more. Supernatants were combined and dried using a vaccuum concentrator at ambient temperature, stored at -80°C, and analyzed within 48 hours.

Metabolite extract samples were reconstituted in HPLC-grade water and analyzed by HPLC (Vanquish Duo UHPLC, Thermo Fisher Scientific) using a hydrophilic interaction chromatography column (XBridge BEH Amide XP Column, 130 Å, 2.5 µm, 2.1 mm × 150 mm, Waters), coupled to a high-resolution orbitrap mass spectrometer (Q Exactive Plus, Thermo Fisher Scientific). MS was performed in both positive and negative mode using a mass resolution of 140,000 at 200 m/z. Data was processed using MAVEN^14^ and corrected for natural isotope abundance using AcuCorr^15^.

### Metabolic flux analysis

^13^C -metabolic flux analysis (^13^C -MFA) was performed using INCA^16^. Briefly, a model of central carbon metabolism, consisting of glycolysis, the pentose phosphate pathway, the TCA cycle, fermentation, and coarse-grained biomass production reactions was constructed, and was fit to ^13^C labeling patterns of metabolites in these pathways. We included constraints based on extracellular flux measurements (glucose consumption, oxygen consumption, ethanol production, glycerol production, and pyruvate production), growth rate measurements, and biomass composition measurements from^13,17,18^. For each condition, the best-fit flux solution was chosen from 200 alternative solutions with randomized initializations (results in Table S6).

### Absolute quantification of NADH

We quantified NADH in two independent ways: via LC-MS and by FLIM. For LC-MS-based quantification, we extracted metabolites from samples of interest as well as a reference sample (prototrophic W303 grown in YNB + 2% glucose) for which absolute quantification of a large number of metabolites, including NADH, has already been performed^19^. We calculated the NADH concentration in the unknown sample by comparing to the reference. For IAA titration, a Michaelis-Menten model was fit to the OCR and LC-MS measurements of NADH concentration:

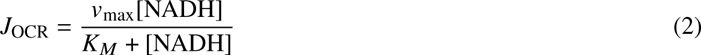

For FLIM-based quantification, we imaged NAD(P)H in live cells as described above. We assume that the molecular brightness of a species is proportional to its fluorescence lifetime^20^. The concentration of NADH is then proportional to intensity *I* and a constant *γ* that depends on experimental parameters such as laser power and detection efficiency:

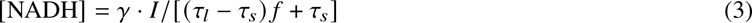

We constructed a calibration curve using standard solutions of NADH, and used this to calculate the NADH concentration in the unknown samples. However, because NADH and NADPH both contribute to the measured fluorescence in cells, a Michaelis-Menten model with an offset was used to relate OCR and NAD(P)H concentrations measured by FLIM:

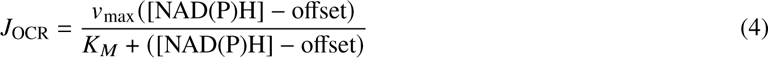

When the fitted value of the offset was subtracted from the measured NAD(P)H concentration, it yielded a curve similar to that obtained from LC-MS measurements of NADH (Fig. S5D). This is consistent with a significant but relatively constant pool of NADPH contributing to the observed intensity.

### Proteomics measurements

Exponentially growing cells were harvested by centrifugation for 2 min at 1500 x g at 4°C. Cells were washed once in ice-cold deionized water, then resuspended in 1 mL ice-cold resuspension buffer (50 mM HEPES pH 7.2 with cOmplete Mini EDTA-free Protease Inhibitor Cocktail (Roche)). The cell suspension was added dropwise to a bath of liquid nitrogen. The same procedure was followed for an equal volume of resuspension buffer containing 4% w/v SDS. Frozen yeast and lysis buffer were added to grinding jars pre-chilled to -196°C, which were then shaken for 3 min at a rate of 15 s^−1^ in a MM400 cryomill (Retsch). Jars were then removed and re-cooled in liquid nitrogen. This procedure was repeated five more times.

Samples were prepared mostly as previously described^21^. Concentrations were determined by reducing agent-compatible Bicinchoninic acid (BCA) assay (Pierce). To reduce disulfide bonds, dithiothreitol (DTT) was added to a final concentration of 5 mM and samples were incubated for 20 min at 60°C. After cooling to room temperature, cysteines were alkylated using N-ethyl maleimide (NEM; final concentration of 20 mM) for 20 min at room temperature. NEM was quenched by an excess of 10 mM DTT. Protein was purified by SP3 precipitation^22^ using magnetic beads (SpeedBead Magnetic Carboxylate, cytiva) at 50% ethanol, then washed three times in 80% ethanol. Protein was digested overnight with 20 ng/µL LysC (Wako) in 2 M guanidine hydrochloride and 10 mM EPPS (pH 8.5) with agitation at 24°C. This was then diluted fourfold with 10 mM EPPS, and an additional 20 ng/µL LysC and 10 ng/µL trypsin (Promega) were added; this was incubated overnight at 37°C. Samples were vacuum-dried, resuspended in 200 mM EPPS (pH 8.0) to a peptide concentration of 1 µg/µL. Labeling of each sample with TMTpro (Thermo Scientific) tags was performed for 2 hr at room temperature at a 5:1 mass ratio of TMTpro to peptide, then quenched with 0.5% hydroxylamine for 30 min at room temperature before combining different conditions. Samples were acidified (to pH < 2) with phosphoric acid and cleared by ultracentrifugation. Supernatants were dried using a vacuum evaporator at RT. The resuspended sample was sonicated for 5 minutes and then fractionated by medium pH reverse-phase HPLC (Zorbax 300Extend C18, 4.6 x 250 mm column, Agilent). The 96 elutions were pooled into 24 fractions by alternating the wells in the plate^23^. Each fraction was dried and resuspended in 100 µL of HPLC water, acidified to pH < 2 with HPLC-grade phosphoric acid, and stage-tipped (C18-tips, Pierce) for desalting^24^. About 2 µg per fraction in 1% formic acid was analyzed in 90 min by LC-MS on an Orbitrap Ascend (Thermo Fisher Scientific) using a Real-Time-Search MS3 method^25^. A quality control sample for cysteine-containing peptides, missed cleavages, labeling efficiency, and channel loading was stage-tipped and analyzed by single-shot LC-MS. Three biological replicate cultures were harvested and prepared as described above for each growth condition. One replicate of each condition was included in each TMTpro experiment; three separate experiments were performed.

### Proteomics data analysis

Mass spectrometry (MS) data analysis was conducted using the Gygi Lab software platform (GFY Core Version 3.8) from Harvard University as previously described^26^. Data in Thermo RAW format were converted to mzXML format, correcting errors in peptide ion charge state and monoisotopic m/z assignments^27^. Monocle software^28^ supported monoisotopic mass detection, and ReAdW.exe was modified to include signal-to-noise ratios during conversion (http://sashimi.svn.sourceforge.net/viewvc/sashimi/). MS2 spectra assignments used the SEQUEST algorithm^29^, searching against databases including the *Saccharomyces cerevisiae* proteome^30^, common contaminants, and reverse protein sequences as decoys. Search parameters included specific ion tolerances and modifications, such as TMTpro tags on lysine and peptide N-termini, and NEM on cysteine. A target-decoy strategy^31^ maintained the false discovery rate of assignments in MS2 spectra below 1%, filtering z-scored spectra and peptide properties by a linear discriminator^27^. Calibration adjusted mass errors in MS1 and MS2 spectra, and peptides were assigned to proteins based on unique matches. The mass spectrometry proteomics data have been deposited to the ProteomeXchange Consortium via the PRIDE^32^ partner repository with the dataset identifier PXD053535.

We considered only those proteins which were measured in all three experiments. Ion abundances were first normalized by the mean across all channels, then by the median across all proteins quantified within a single channel. The group abundances reported here represent the sum of these normalized abundances of each protein in the group. Members of each group are listed in Table S3. Localization data was obtained from Uniprot. For organelle-level analyses, we included only those proteins which were localized to a single organelle.

## Acknowledgements

We thank Vladimir Denic, Matthias Heinemann, Andrew Murray, Yong Hyun Song, and Xingbo Yang for valuable discussions; Laura Bagamery and Piyush Nanda for the kind gift of strains and plasmids; Joshua Rabinowitz for guidance on metabolomics; Dongtao Cui and Anthony Lowe of the Laukien-Purcell Instrumentation Center for guidance on NMR measurements; and Gloria Ha and Michael van der Naald for a critical reading of the manuscript. This work was supported by National Science Foundation award MCB-2052305 to D.J.N.; grants from the National Institutes of Health (under award number R35GM128813), the Simons Foundation, and the Princeton Catalysis Initiative to M.W.; and a grant from the National Institutes of Health (under award number R35GM143127) to J.O.P.

## Author Contributions

E.A., J.O.P., M.W., and D.J.N. designed the study. E.A., F.C.K., R.C.L., and C.K.K. performed experiments. E.A., F.C.K., R.C.L., and Y.S. analyzed data. E.A. and D.J.N. wrote the paper with input from all authors.

## Data and code availability

Source data are available in supplementary tables. Proteomics data are available in the PRIDE partner repository with the dataset identifier PXD053535. The library pyTCSPC, developed for analyzing the FLIM experiments in this study, is available at https://www.github.com/easunarunachalam/pyTCSPC. Sample code for mitochondrial network analysis is available at https://www.github.com/easunarunachalam/SADmodel.

