## Supplementary Text and Figures for "Robustness of mitochondrial biogenesis and respiration explain aerobic glycolysis"

- Table S1: Strains used in this work
- Table S2: Protein abundances from TMT experiments
- Table S3: Proteins included in candidate groups for regression against respiration rate
- Table S4: Extracellular flux measurements and biomass fluxes
- Table S5: Labeling data from isotope tracing experiments
- Table S6: Inferred fluxes from $^{13}\text{C}$-MFA of glucose-limited cultures
- Table S7: Kinetic parameters for major hexose transporters
- Supplementary Note 1: Mathematical model of mitochondrial growth
- Supplementary Note 2: Michaelis-Menten model of glucose consumption rate
- Supplementary Note 3: Model of NADH production and consumption

#### Supplementary Note 1: Mathematical model of mitochondrial growth

Let us denote the mitochondrial volume in a single cell over the course of a cell cycle by $v(t)$. At the beginning of the cell cycle, the mitochondrial volume is $v(0) \equiv v_i$; at the end of the cell cycle (duration $\tau$), the volume is $v(\tau) \equiv v_f$. Let $\Delta v(t)$ denote the difference between the initial volume and the volume at some point $t$. Then, we have:

$$v(t) = v_i + \Delta v(t) \quad (\text{S1})$$

with $\Delta v(0) = 0$ by construction. Over the course of the cell cycle, the mitochondrial volume increases by some factor $f \equiv v_f/v_i$:

$$v(\tau) = v_i + \Delta v(\tau) = f v_i$$

Then,

$$v(t) = \frac{\Delta v(\tau)}{f-1} + \Delta v(t)$$

Therefore, the mean volume of the cell over the course of the cell cycle is:

$$\bar{v} = \frac{1}{\tau} \int_0^\tau v(t) dt = \frac{1}{\tau} \int_0^\tau \left( \frac{\Delta v(\tau)}{f-1} + \Delta v(t) \right) dt = \frac{\Delta v(\tau)}{f-1} + \overline{\Delta v}$$

where $\bar{\cdot}$ indicates an average over the cell cycle.

Mitochondrial growth occurs in two stages: during G1 (duration $\tau_g$) the mitogenesis rate is $r_g$, and during budding (duration $\tau_b$) the mitogenesis rate is $r_b$ (Fig. S6A). (Note that the length of the cell cycle is $\tau = \tau_g + \tau_b$.) The cell-cycle-averaged mitochondrial biogenesis rate $\bar{r}$ is the harmonic mean of these rates:

$$\bar{r} = \frac{r_g \tau_g + r_b \tau_b}{\tau_g + \tau_b} \quad (\text{S2})$$

The volume change over time is given by:

$$\Delta v(t) = \begin{cases} r_g t & 0 \leq t < \tau_g \\ r_g \tau_g + r_b (t - \tau_g) & \tau_g \leq t < \tau_g + \tau_b \end{cases}$$

and its average value is:

$$\overline{\Delta v} = \frac{1}{2} (r_g \tau_g + \bar{r} \tau_b) \quad (\text{S3})$$

Thus, the average mitochondrial volume over the course of the cell cycle is:

$$\bar{v} = \bar{r}\tau \left( \frac{1}{2} + \frac{1}{f-1} \right) - \frac{1}{2} (\bar{r} - r_g) \tau_g$$

Using data from the conditions for which we directly measured mitochondrial biogenesis rates, we find that the average magnitude of the second term does not exceed 5% of that of the first term. Hence, the average mitochondrial volume in a single cell is approximately proportional to the average biogenesis rate $\bar{r}$ and division time $\tau$:

$$\bar{v} \approx \bar{r}\tau \left( \frac{1}{2} + \frac{1}{f-1} \right) \quad (\text{S4})$$

The average mitochondrial volume across a population depends on the mitochondrial biogenesis rate, cell cycle time, and final-to-initial volume ratio in individual cells. The simplest assumption is that each of these quantities is an independent random variable. Denoting an average over the population by $\langle \cdot \rangle$, we have that the population-average volume $\langle V \rangle$ is then:

$$\langle V \rangle \approx \langle \bar{r} \rangle \langle T \rangle \left( \frac{1}{2} + \frac{1}{\langle f \rangle - 1} \right) \quad (\text{S5})$$

where $\langle T \rangle$ is the bulk doubling time.

### Supplementary Note 2: Michaelis-Menten model of glucose consumption rate

The relative abundance (RA) of each major hexose transporter in each of the glucose-limited cultures was measured by proteomics. The Michaelis constant $K_M$ and maximal velocity $v_{\max}$ for each hexose transporter were obtained from previous work (Table S7):

| Hexose transporter $i$ | $K_M$ (mM) | $v_{\max}$ (nmol min<sup>-1</sup> mg<sup>-1</sup>) | Reference |
| --- | --- | --- | --- |
| *HXT1* | 129 ± 9 | 50.9 ± 3.7 | [1] |
| *HXT2* | 2.8 ± 0.1 | 35.3 ± 1.3 | [1] |
| *HXT3* | 34.2 ± 3.2 | 18.5 ± 2.0 | [1] |
| *HXT4* | 6.2 ± 0.3 | 12.0 ± 0.9 | [1] |
| *HXT5* | 10 ± 1 | 150 ± 20 | [2] |
| *HXT6* | 1.4 ± 0.1 | 11.4 ± 0.5 | [1] |
| *HXT7* | 1.9 ± 0.1 | 11.7 ± 0.3 | [1] |

Table S7: Kinetic parameters for major hexose transporters.

The relative glucose consumption rate (GCR) for transporter $i$ was calculated as:

$$\widehat{\text{GCR}}^{(i)} = \frac{\text{RA}^{(i)} v_{\max}^{(i)} [\text{glucose}]}{K_M^{(i)} + [\text{glucose}]} \quad (\text{S6})$$

The total glucose consumption rate (GCR) for condition $j$ was calculated as:

$$\widehat{\text{GCR}} = \sum_i \widehat{\text{GCR}}^{(i)} \quad (\text{S7})$$

using point estimates for the kinetic parameters. This was normalized to the median value across different conditions.

### 701 Supplementary Note 3: Model of NADH production and consumption

Reducing equivalents (REs) are produced primarily by glycolysis and the tricarboxylic acid (TCA) cycle. The rates of RE production by these pathways over the series of glucose-limited cultures were calculated by $^{13}\text{C}$-MFA. These rates were well-predicted by glucose consumption rate (GCR) using simple empirical models: NADH production by glyceraldehyde-3-dehydrogenase (GAPDH) was fit to a linear model (Fig. S8A) and NADH/FADH<sub>2</sub> production by the TCA cycle (pyruvate dehydrogenase, isocitrate dehydrogenase, α-ketoglutarate dehydrogenase, succinate dehydrogenase, and malate dehydrogenase) was fit to a quadratic model (Fig. S8B):

$$\begin{aligned} J_{\text{RE prod, glycolysis}}(\text{GCR}) &= \alpha_{\text{GAPDH}} + \beta_{\text{GAPDH}} \text{GCR} \\ J_{\text{RE prod, TCA}}(\text{GCR}) &= \alpha_{\text{TCA}} + \beta_{\text{TCA}} \text{GCR} + \gamma_{\text{TCA}} \text{GCR}^2 \end{aligned} \quad (\text{S8})$$

where $\alpha$., $\beta$., and $\gamma$. indicate regression coefficients. The total production of reducing equivalents was thus predicted using the model for the total glucose consumption rate (discussed in Supplementary Note 2):

$$\hat{J}_{\text{RE prod}}(\widehat{\text{GCR}}) = J_{\text{RE prod, glycolysis}}(\widehat{\text{GCR}}) + J_{\text{RE prod, TCA}}(\widehat{\text{GCR}}) \quad (\text{S9})$$

At steady state, the total rate of RE production is matched by the total rate of RE consumption, and the primary RE-consuming reactions are respiration and fermentation:

$$\begin{aligned} J_{\text{RE prod}} &= J_{\text{RE cons}} = J_{\text{RE cons, resp}} + J_{\text{RE cons, ferm}} \\ &= 2 \times (J_{\text{oxygen cons}}) + 1 \times (J_{\text{ethanol prod}} + J_{\text{glycerol prod}}) \end{aligned} \quad (\text{S10})$$

The measured total RE production reported in Fig. 6B is therefore calculated from measured oxygen consumption rates and ethanol and glycerol production rates.

The rate of RE consumption by respiration was predicted using the relationships between doubling time, mitochondrial volume, and respiration rate (i.e. the SAD model). The linear relationship between OCR and mean single-cell mitochondrial volume $V$, i.e. $\text{OCR} = \alpha_{\text{resp}} + \beta_{\text{resp}} V$ where $\alpha_{\text{resp}}, \beta_{\text{resp}}$ are regression coefficients, is illustrated in Fig. 2B. The cell-cycle-averaged mitochondrial biogenesis rate $\bar{r}^*$ in glucose-limited cultures was estimated from a regression of mean single-cell mitochondrial volume against population doubling time (Fig. S8C). The OCR (and thus respiratory NADH consumption rate) under each condition was therefore estimated using only the population doubling time under that condition:

$$\begin{aligned} \widehat{\text{OCR}}(\langle T \rangle) &\approx \alpha_{\text{resp}} + \beta_{\text{resp}} \left( \bar{r}^*(\langle T \rangle) \left( \frac{1}{2} + \frac{1}{\langle v_f/v_i \rangle - 1} \right) \right) \\ \hat{J}_{\text{RE cons, resp}}(\langle T \rangle) &= 2 \times \widehat{\text{OCR}}(\langle T \rangle) \end{aligned} \quad (\text{S11})$$

The rate of RE consumption by fermentation was taken as the difference between the total rate of RE production and the rate of RE consumption by respiration:

$$\hat{J}_{\text{RE cons, ferm}}(\widehat{\text{GCR}}, \langle T \rangle) = \hat{J}_{\text{RE prod}}(\widehat{\text{GCR}}) - \hat{J}_{\text{RE cons, resp}}(\langle T \rangle) \quad (\text{S12})$$

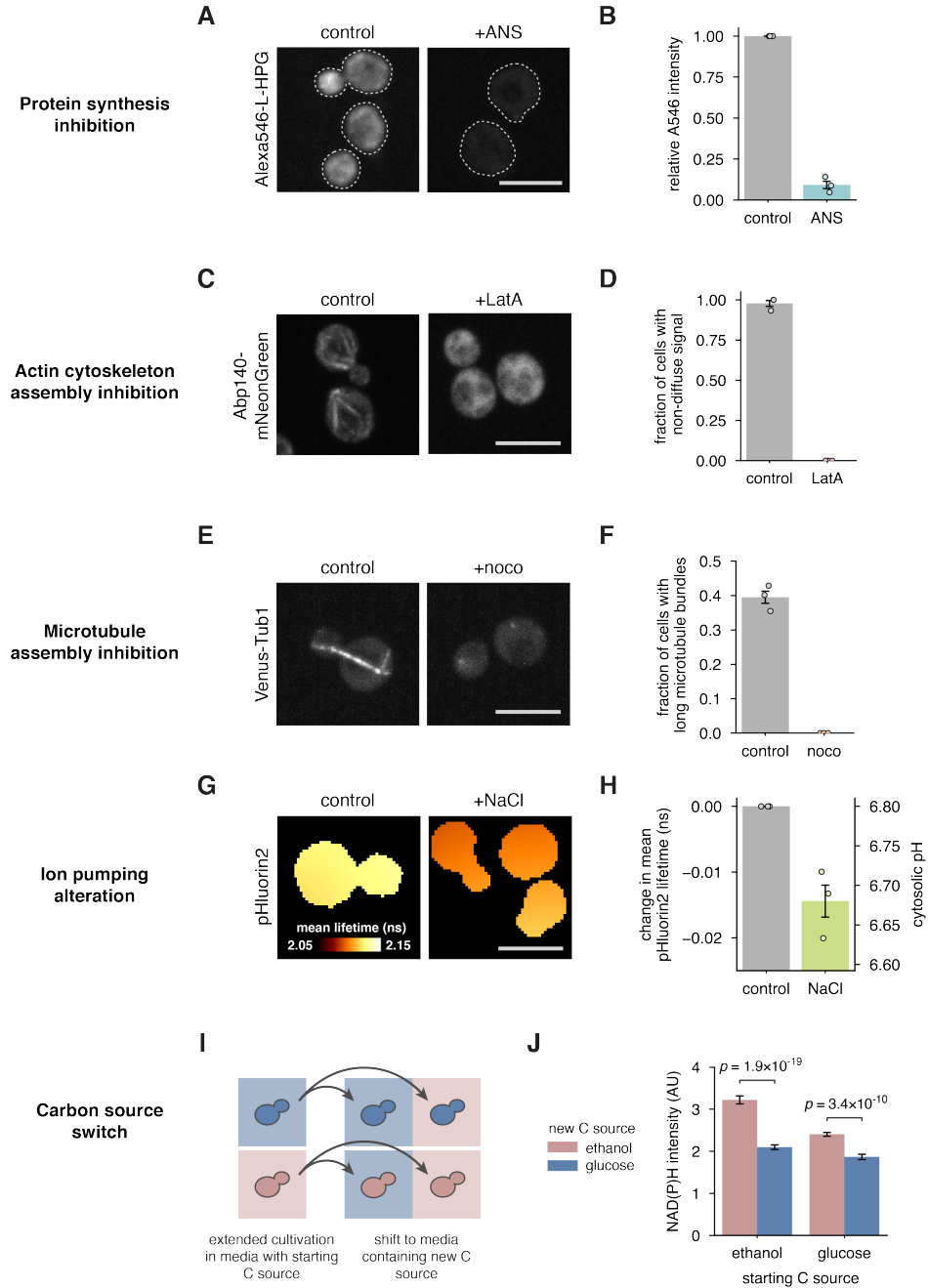

**Figure S1: Verification of ATP demand and nutrient supply perturbations.** (A) Alexa546-labeled L-HPG incorporated into newly synthesized proteins in control and 300 μM anisomycin (ANS)-treated cells. Cell boundaries are indicated by dashed lines for visibility. (B) Relative intensity of Alexa546 fluorescence in control and ANS-treated cells. (C) Actin network morphology visualized using actin-binding protein Abp140-mNeonGreen in control and 200 μM Latrunculin A (LatA)-treated cells. (D) Fraction of cells with clearly visible subcellular structures, i.e. non-diffuse signal, under each condition. (E) Tubulin association visualized using Venus-Tub1 in control and 66 μM nocodazole (noco)-treated cells. (F) Fraction of cells with long microtubule bundles visible under each condition. (G) pH changes associated with 200 mM NaCl treatment visualized by spatially averaged fluorescence lifetime images of the biosensor pHluorin2. (H) Changes in mean lifetime and pH under each condition. (I) Cells grown for an extended period in one carbon source were shifted to another carbon source before metabolic measurements. (J) NAD(P)H autofluorescence changes significantly upon shifting ethanol-grown cells to glucose, and vice versa. In (A, C, E, G) scale bars are 5 μm. In (B, D, F, H) data are shown as mean ± s.e.m. ($n = 3$ biological replicates). In (J) data are shown as mean ± s.e.m. of $n \geq 59$ images across 3 (starting from glucose) or 4 (starting from ethanol) biological replicates.

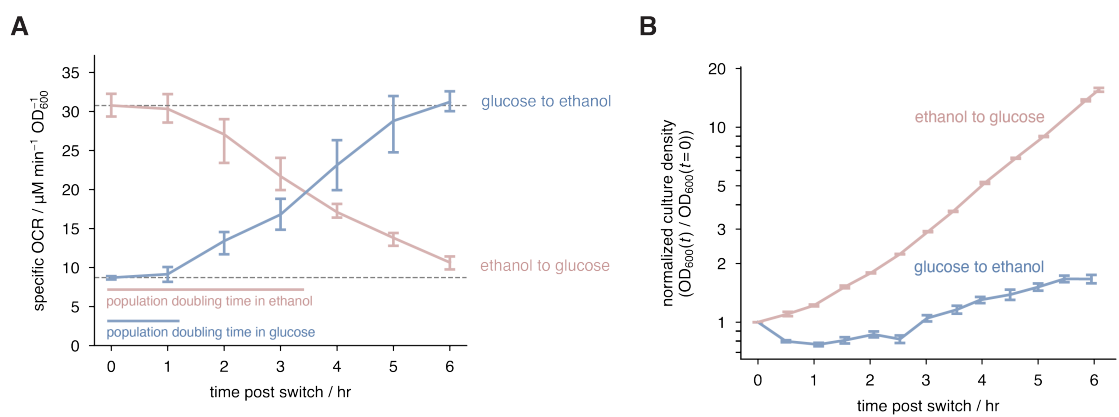

**Figure S2: OCR adapts on the timescale of cell growth and division following nutrient shifts.** Dynamics of (A) OCR and (B) culture density following nutrient shifts. Shifts from glucose to ethanol are shown in blue, and shifts from ethanol to glucose are shown in red. Data are shown as mean ± s.e.m. ($n \geq 3$ biological replicates for OCR and $n = 3$ biological replicates for culture density). In (A) grey dashed lines indicate the initial OCR of ethanol- and glucose-grown cells, and scale bars indicate population doubling times of exponential-phase ethanol- and glucose-grown cells.

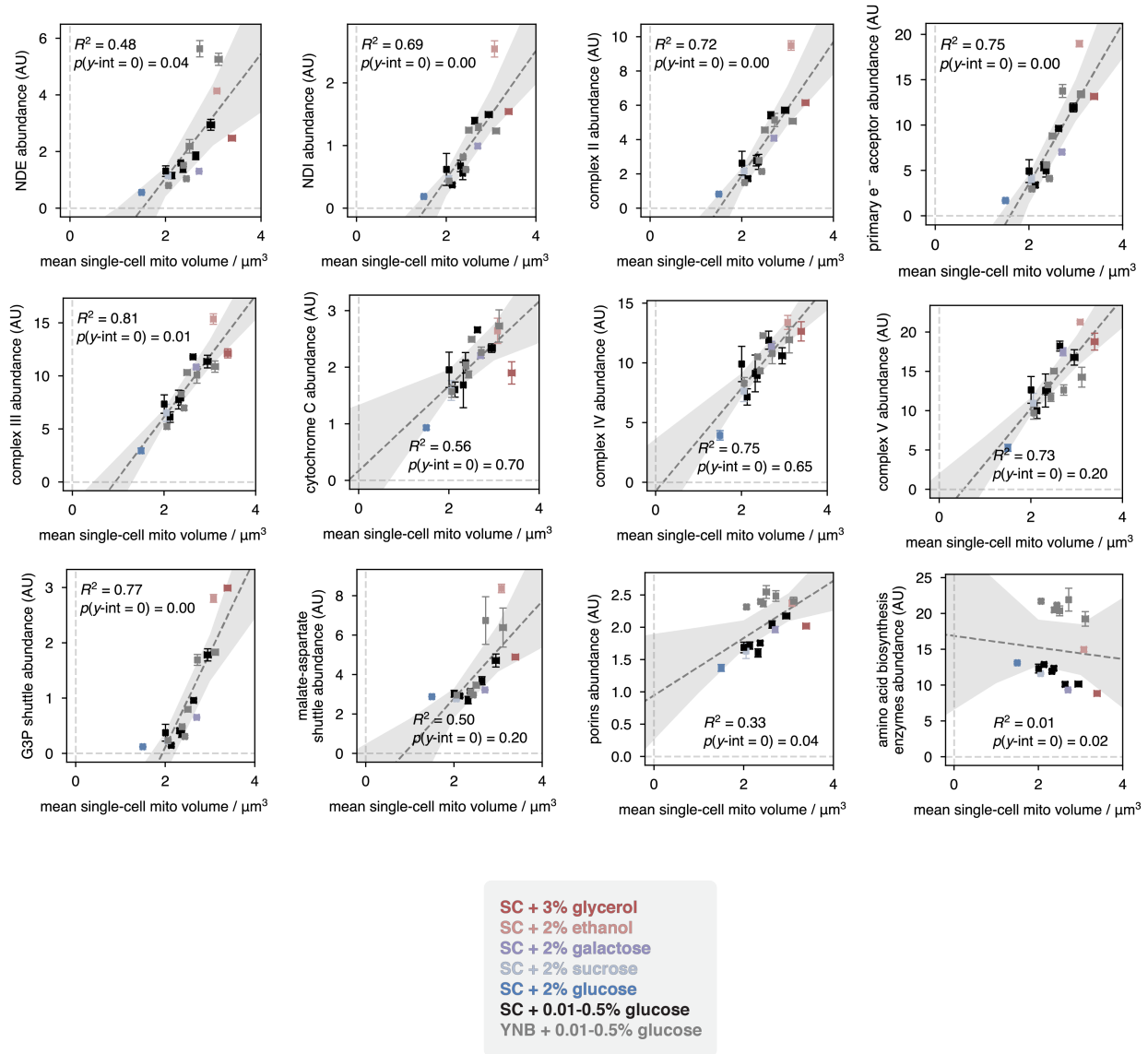

**Figure S3: Relationship between abundance of mitochondrial proteins and mitochondrial volume measured by microscopy.** Linear regressions of selected respiration-related and other mitochondrial proteins against mitochondrial volume. Data are shown as mean ± s.e.m. ($n \geq 3$ biological replicates for mitochondrial volume measurements, and $n = 3$ biological replicates for proteomics); shaded regions reflect 95% confidence intervals for linear regression. $p$-values indicate the probability of observing the associated $y$-intercept, or one more extreme, given the null hypothesis that the $y$-intercept is zero, and were estimated by bootstrapping.

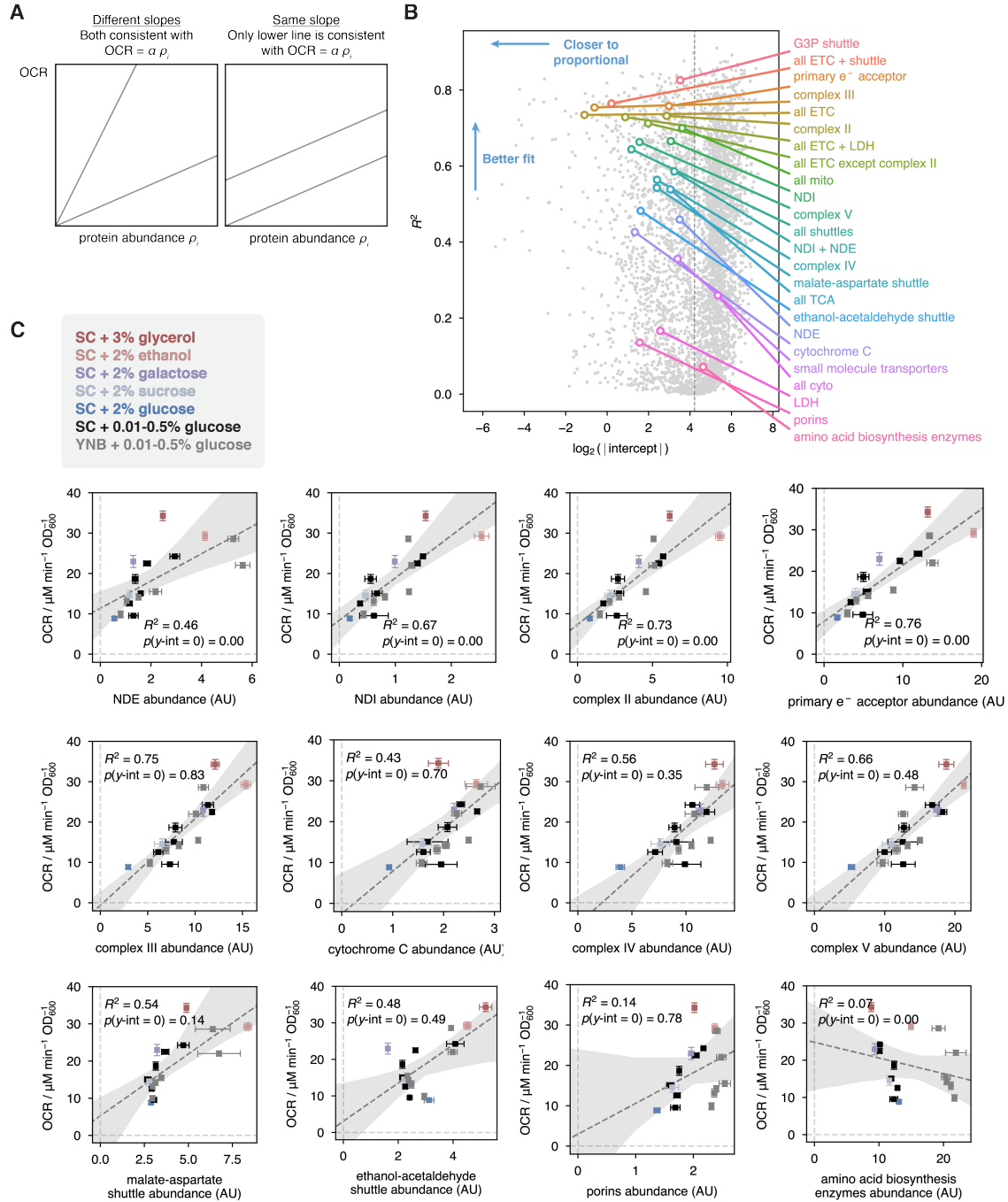

**Figure S4: Relationship between OCR and abundance of candidate protein groups.** (A) Schematic of two different types of relationships: the abundance of a group may be proportional to OCR but with different slopes (left) or may be linearly related to OCR but with different y-intercepts (right). Only those protein groups with a y-intercept of zero can plausibly control OCR. (B) $R^2$ vs. y-intercept for OCR-protein abundance regressions. The dashed vertical reference line indicates the average y-value of all data points used in the regression, i.e. the mean OCR across different conditions. All quantified proteins are shown as gray dots, while candidate groups are shown in color. (C) Regressions of OCR against abundance of selected protein groups. Data are shown as mean ± s.e.m. ($n \geq 3$ biological replicates for OCR, $n = 3$ biological replicates for proteomics); shaded regions reflect 95% confidence intervals for linear regression. $p$-values indicate the probability of observing the associated y-intercept, or one more extreme, given the null hypothesis that the y-intercept is zero, and were estimated by bootstrapping.

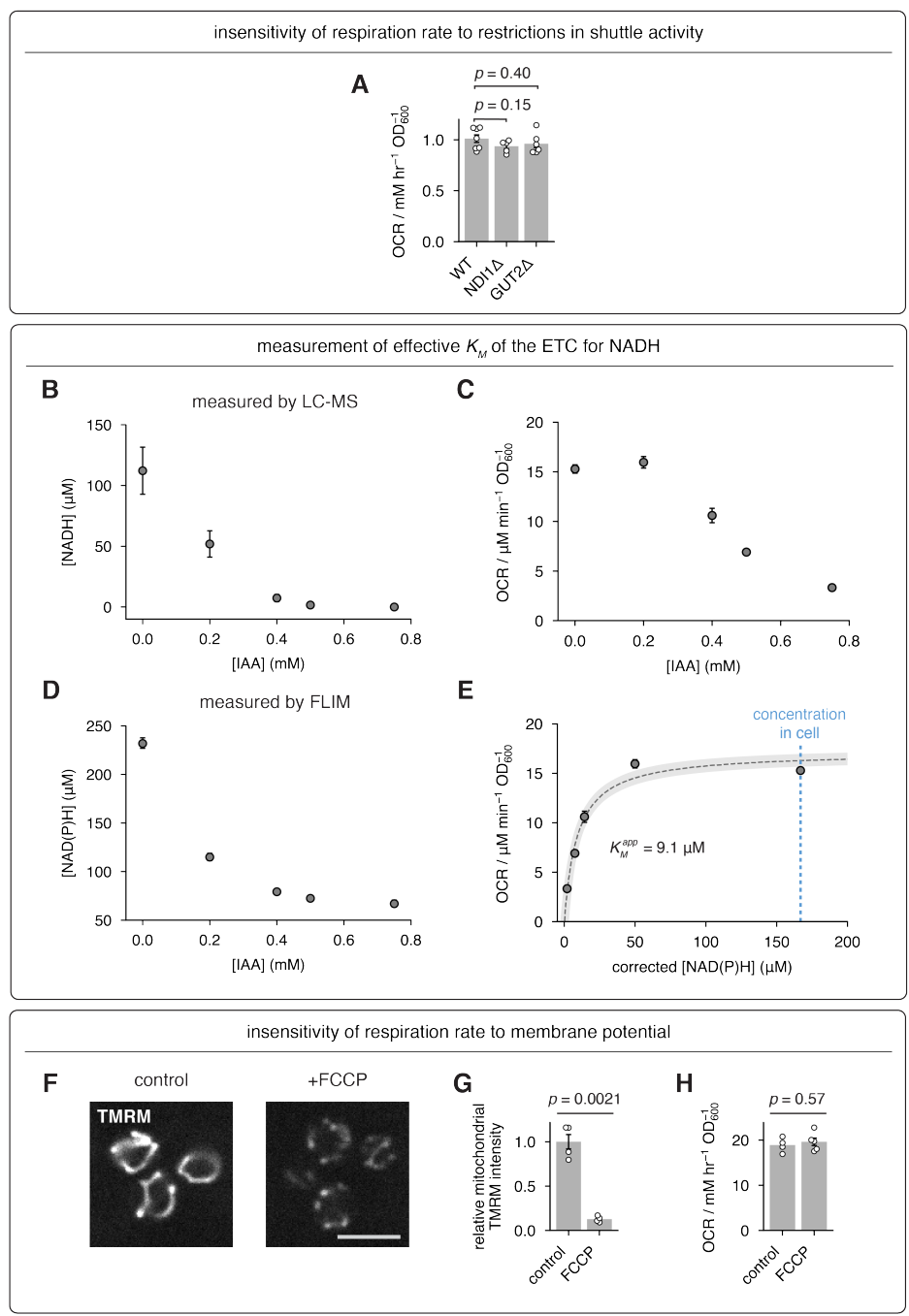

**Figure S5: Respiration is saturated by NADH and is insensitive to changes in mitochondrial membrane potential.** (A) Respiration rate is insensitive to knockouts of *ND1* and *GUT2* ($n \geq 5$ biological replicates). Data are shown as mean ± s.e.m. and $p$-values are obtained by Welch's $t$-test. (B) NADH concentration measured by LC-MS as a function of IAA concentration ($n = 3$ biological replicates). (C) Oxygen consumption rate as a function of the concentration of the GAPDH inhibitor IAA ($n \geq 4$ biological replicates). (D) NAD(P)H concentration measured by FLIM as a function of IAA concentration ($n = 4$ biological replicates). (E) Oxygen consumption rate as a function of NAD(P)H concentration. Dashed line is the best fit to a Michaelis-Menten model with concentration offset (see methods), and the $x$-axis reflects the measured NAD(P)H concentration less the fitted offset. The shaded region reflects the 95% confidence interval from bootstrapping. In (B)-(E) data are shown as mean ± s.e.m. (F) Mitochondrial membrane potential in glucose-grown cells, visualized using TMRM. Left, untreated cells; right, cells treated with 200 nM carbonyl cyanide-*p*-trifluoromethoxyphenylhydrazone (FCCP). Scale bar is 5 μm. (G) FCCP treatment significantly decreases membrane potential ($n = 4$ biological replicates), but (H) does not significantly alter OCR ($n \geq 4$ biological replicates). Data are shown as mean ± s.e.m. and $p$-values are obtained by Welch's $t$-test.

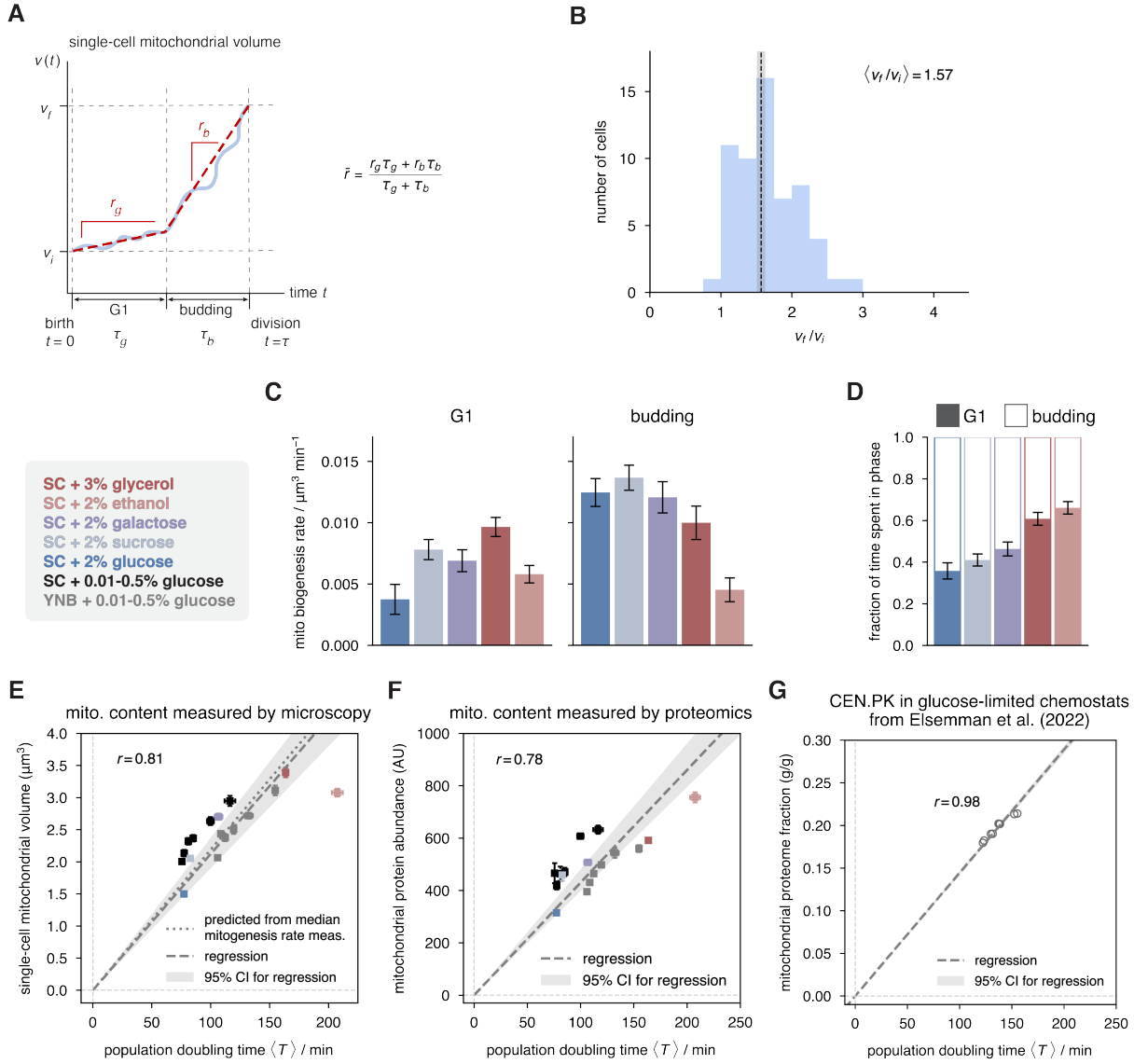

**Figure S6: Details of mitochondrial biogenesis rate measurement and modeling.** (A) Schematic trajectory of mitochondrial volume within a single cell over the course of a cell cycle. Over the course of the G1 phase of the cell cycle (of duration $T_g$), mitochondrial volume accumulates at a rate $r_g$. Over the course of budding (duration $T_b$) mitochondrial volume accumulates at a rate $r_b$. The mean rate of mitochondrial biogenesis over the cell cycle, $\bar{r}$, is given by the harmonic mean of $r_g$ and $r_b$. (B) Ratio of mitochondrial volume at the end of the cell cycle, $v_f$, to that at the beginning of the cell cycle, $v_i$, for $n = 59$ cells across $n = 4$ independent experiments. The median value is $\langle v_f/v_i \rangle = 1.57$. (C) Measured mean mitochondrial biogenesis rates during G1 and budding across different carbon sources. (D) Fraction of time spent in G1 vs. budding for cells grown with each carbon source. In (C) and (D) data are shown as mean ± s.e.m. from $n \geq 151$ cells, across G1 and budding from $n = 3$ biological replicates, for each carbon source. (E)-(F) Consistent with the SAD model, mitochondrial content across a range of conditions is approximately proportional to division time, as measured by (E) microscopy ($n \geq 3$ biological replicates), and (F) proteomics ($n = 3$ biological replicates). Data are shown as mean ± s.e.m. (G) Mitochondrial proteome fraction measurements in another *S. cerevisiae* strain, cultivated in high-dilution-rate glucose-limited chemostats, are proportional to doubling time. Data are from Elsemman *et al.* [3]; each circle represents an independent experiment. In (E)-(G) dashed lines indicate linear regressions, and shaded regions indicate the 95% confidence intervals from bootstrap sampling.

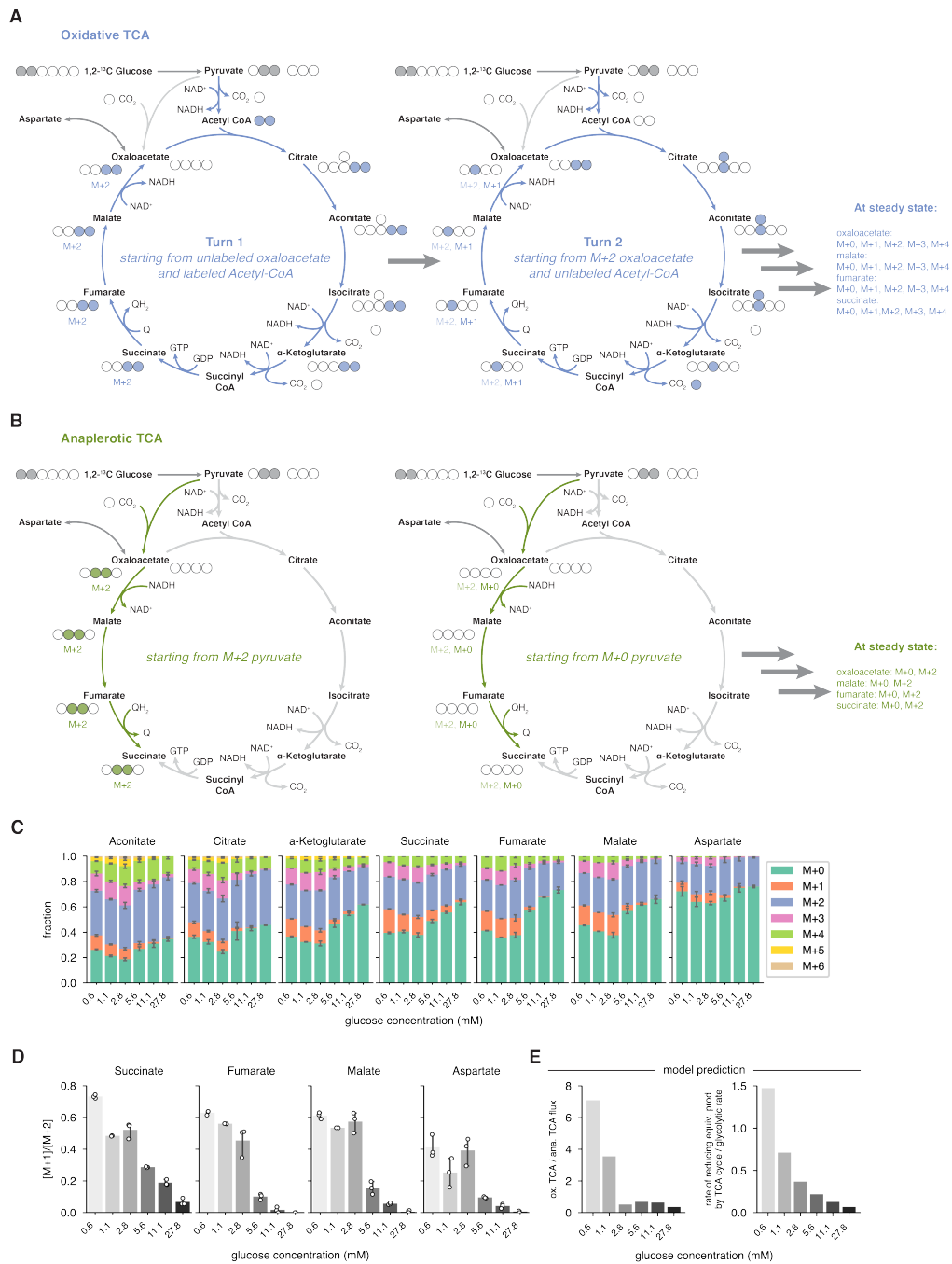

**Figure S7: The transition from oxidative to anaplerotic TCA metabolism does not coincide with the onset of ethanol production.** (A) When supplied with 1,2-$^{13}\text{C}$ glucose, oxidative TCA cycle turning produces a mixture of isotopomers of 4-carbon intermediates, while (B) anaplerosis produces only M+0 and M+2 isotopomers. $^{12}\text{C}$ atoms are indicated by unfilled circles, and $^{13}\text{C}$ atoms by filled circles. (C) Mass isotopomer distributions of TCA cycle intermediates for cells cultivated in media with different glucose concentrations. Data are shown as mean ± s.e.m. The presence of M+1 and M+3 isotopomers of 4-carbon intermediates, and the qualitative similarity of labeling patterns across different glucose concentrations, indicates that the TCA cycle turns fully even under some conditions where glucose is abundant and the rate of fermentation is high. (D) The ratio of [M+1] to [M+2] isotopomers of 4-carbon TCA cycle intermediates is a model-independent indicator of relative fluxes through the oxidative and anaplerotic arms of the TCA cycle, and decreases with increasing glucose concentration. (E) The ratio of oxidative TCA flux to anaplerotic TCA flux (succinate dehydrogenase to pyruvate carboxylase flux) obtained from the fitted model similarly declines with increasing glucose concentration. The rate of reducing equivalent production by the TCA cycle relative to glycolytic rate follows a similar trend.

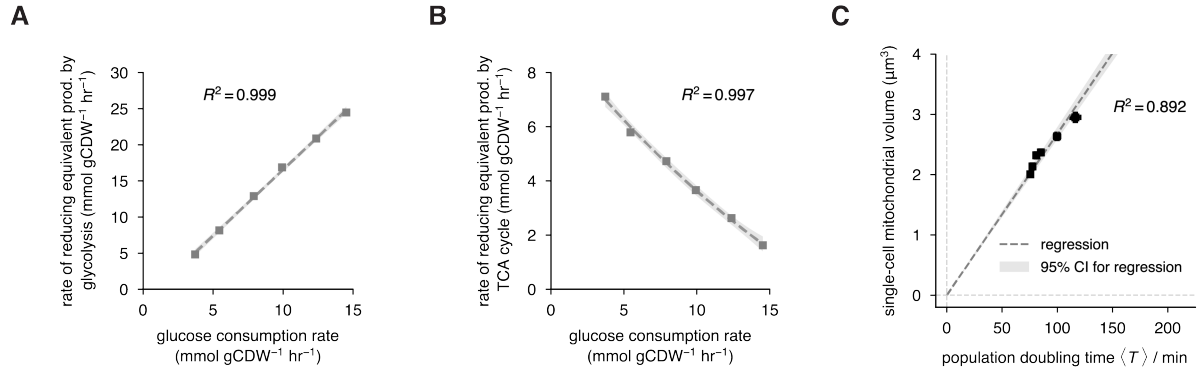

**Figure S8: Components of the NADH production model.** (A) The rate of NADH production by glyceraldehyde-3-phosphate dehydrogenase (GAPDH) as a function of glucose consumption rate, as predicted by <sup>13</sup>C -MFA. The dashed line indicates a linear fit. (B) The rate of NADH and FADH<sub>2</sub> production by the TCA cycle as a function of glucose consumption rate, as predicted by <sup>13</sup>C -MFA. The dashed line indicates a quadratic fit. Shaded regions indicate 95% confidence intervals for the fit. (C) A fit of mean single-cell mitochondrial volume to population doubling time for glucose-limited cultures enables estimation of cell-cycle-averaged mitochondrial biogenesis rate under these conditions. Shaded regions indicate 95% confidence intervals from bootstrap sampling.
